# Real time observation of chaperone-modulated talin mechanics with single molecule resolution

**DOI:** 10.1101/2021.04.27.441571

**Authors:** Soham Chakraborty, Deep Chaudhuri, Souradeep Banerjee, Madhu Bhatt, Shubhasis Haldar

**Author notes:** contributed equally to this work (Soham Chakraborty, Deep Chaudhuri, Souradeep Banerjee).

## Abstract

Recent single-molecule studies have recognized talin as a mechanosensitive hub in focal adhesion, where its function is strongly regulated by mechanical force. For instance, at low force (below 5 pN), folded talin binds RIAM for integrin activation; whereas at high force (above 5 pN), it unfolds to activate vinculin binding for focal adhesion stabilization. Being a cytoplasmic protein, talin might interact with several cytosolic chaperones: however, the role of chaperones in talin mechanics is unknown.

To address this question, we investigated the force response of a mechanically stable talin domain with a set of well-known holdase (DnaJ, DnaK, Hsp70, and Hsp40) and foldase (DnaKJE, DsbA) chaperones, using single-molecule magnetic tweezers. Our findings demonstrate that chaperone could affect adhesion proteins stability by changing their folding mechanics; while holdase chaperones reduce their unfolding force to ∼6 pN, foldase chaperones shift it up to ∼15 pN. Since talin is mechanically synced within 2 pN force ranges, these changes are significant in cellular condition. Furthermore, we determined the fundamental mechanism of this altered mechanical stability, where chaperones directly reshape their energy landscape: unfoldase chaperone (DnaK) decreases the unfolding barrier height from 26.8 to 21.7 k_B_T, while foldase chaperone (DsbA) increases it to 33.5 k_B_T. We reconciled our observations with eukaryotic Hsp70 and Hsp40 chaperones and observed their similar function of decreasing the talin unfolding barrier to 23.1 k_B_T. The quantitative mapping of this chaperone-induced talin folding landscape directly illustrates that chaperones perturb the adhesion protein stability under physiological force, thereby influencing their force-dependent interactions and adhesion dynamics.

## Introduction

Cells interpret and respond to complex mechanical environments through focal adhesion (FA) mediated responses which help to transduce mechanical cues into biochemical signals, supporting diverse biological processes such as, cell development, migration, and proliferation.^1^ These mechanical communications, through the large adhesion complex, are tightly regulated by the physical linkages and elasticity of mechanosensing proteins including intermediate filaments and multidomain proteins.^2–4^ Talin is a key mechanosensitive scaffold protein which connects transmembrane integrin to actin cytoskeleton, and mechanically tunes diverse interactions by force transmission.^2–5^ Talin rod segment (R1-R13) exhibits conformational changes and hierarchical unfolding of each rod domains under increasing mechanical tension.^6–8^ Among them, R3 is the least stable domain that not only unfolds at ∼5 pN but also exhibits equilibrium folding-unfolding transitions on a sub-second timescale by a single actomyosin contraction.^2,8,9^ This force-dependent folding dynamics allow them to tune their interactome profile: such as, Rap1-interacting adapter molecule (RIAM) binds to folded R3 domain and protects from mechanical unfolding, whereas force above 5 pN results in stepwise unfolding of the domain with subsequent vinculin binding. This mutual exclusive interaction allows talin to acts as a mechanochemical switch, shifting from an initial integrin activator to the active mediator of adhesion maturation and stabilization.^2,10^ Interestingly molecular chaperones, involved in various folding related process starting from protein translation to degradation, could play a pivotal role in FA stabilization by colocalizing with different cytoskeletal and adhesion proteins.^11–13,13–17^ Since Hsp70/Hsp40 are the ubiquitous cytoplasmic chaperones, it interacts with different cytosolic proteins and talin being a critical mechanosensitive cytosolic protein, possess a high probability to interact with the chaperone complex. However, there is no direct evidence on how these chaperones could mechanically influence its interaction profiles during the adhesion maturation process.

To address this question, single-molecule magnetic tweezers have been used, which executes both force ramp and force clamp methodologies.^18^ The force ramp methodology, by increasing or decreasing the force at a constant rate, probes unfolding and refolding events of a protein; while applying the force clamp method, a constant force can be applied to the protein, allowing to detect their thermodynamics and kinetic properties under equilibrium condition. Owing to the advantage of wide force regime of 0-120 pN with sub-pN resolution, we are able to observe chaperone interactions both at folded and unfolded states, independently. Additionally, the flow chamber allows us to introduce chaperones, individually or in combination, to probe their effect on folding dynamics of a single protein in real time. Finally, implementing this methodology, the force can be specifically applied on the client protein while the chaperones remain unperturbed.

Here, we systematically investigated the chaperone effects on the mechanical stability of talin R3 domain, with a set of model chaperones: holdase, foldase and mechanically neutral. Our result showed that DnaK as a holdase chaperone mechanically weakens the R3-IVVI domain, reducing its unfolding force from 10 to 6 pN; whereas oxidoreductase enzyme DsbA as a foldase chaperone increases its stability by increasing the force to 15 pN. However, the mechanically neutral DnaKJE chaperone complex does not exhibit any additional effect on unfolding and refolding force of the substrate. To generalize our hypothesis, we further studied the effect of Hsp70 and Hsp40, the eukaryotic homologues of DnaK and DnaJ system, on the talin mechanical stability. Interestingly, we observed that Hsp70 and Hsp40, very similar to DnaK and DnaJ, function as unfoldases, decreasing the unfolding force of talin from 10.7 pN to 7.7 and 8.3 pN, respectively. Additionally, from the force-induced equilibrium condition, we explicitly demonstrate the underlying physical mechanism of this altered mechanical response by illustrating the quantitative mapping of chaperone-induced folding landscape. We observed chaperones could reshape the mechanical folding landscape of client proteins by changing the height of their free energy barrier without significantly affecting the transition state distance. For example, DnaJ or DnaK stabilizes the unfolded state by decreasing the unfolding barrier height, while DsbA stabilizes the folded state of protein by tilting the energy landscape towards the opposite pathway. We explored our data by Bell-like equation to resolve the quantitative description of the chaperone-modulated free energy landscape, which has not been reported before. From a broader viewpoint, this mechanism may have generic mechanistic insight of how talin response and their binding kinetics with other interactors under force, could be affected by chaperone-altered stability. Overall, our result expanded the canonical function of molecular chaperones and illustrated their novel role in modulating the stability of mechanically stretched protein substrates.

## Results

### Single molecule folding dynamics of Talin R3-IVVI domain

Due to the presence of four threonine residues at its hydrophobic core, R3 domain is the mechanically weakest domain in talin. It unfolds earlier than any other rod domains and exhibits the equilibrium folding dynamics at ∼4-6 pN.^8,19^ Hence, we used a mechanically stable version of R3 domain by substituting its four threonine residues with amino acids containing larger hydrophobic residue-isoleucine and valine at 809, 833, 867 and 901 position, referred as R3-IVVI domain. Both the wild type R3 and R3-IVVI domains have been extensively studied and observed to share similar function, however, R3-IVVI exhibits amplified mechanical signature. For example, both of them bind vinculin at unfolded state but wild type R3 domain unfolds at 5 pN, whereas R3-IVVI unfolds at 9 pN.^10,20,21^ Since both of them exhibit similar functional property, we used R3-IVVI domain as our substrate.

The real time folding-unfolding dynamics of R3-IVVI domain, at single molecular resolution, has been observed using real time magnetic tweezers technology. This technology allowed the application of sub-pN level of force to understand the folding dynamics of the protein at equilibrium. This single-molecule force spectroscopy technology provides two ways for experimentation-force ramp and force clamp techniques. The talin construct is covalently attached to glass surface, using the HaloTag covalent chemistry and the C-terminal Avi-tag was biotinylated to bind the paramagnetic beads through biotin-streptavidin chemistry (Fig. 1A). Force is applied on the paramagnetic bead with a pair of permanent magnets, attached to a voice coil actuator, which controls the amount of force as an inverse function of the distance between permanent magnet and the paramagnetic beads.^22^ The detailed force calibration is provided in the supplemental information (Supplementary Figure 14).

**Figure 1:**
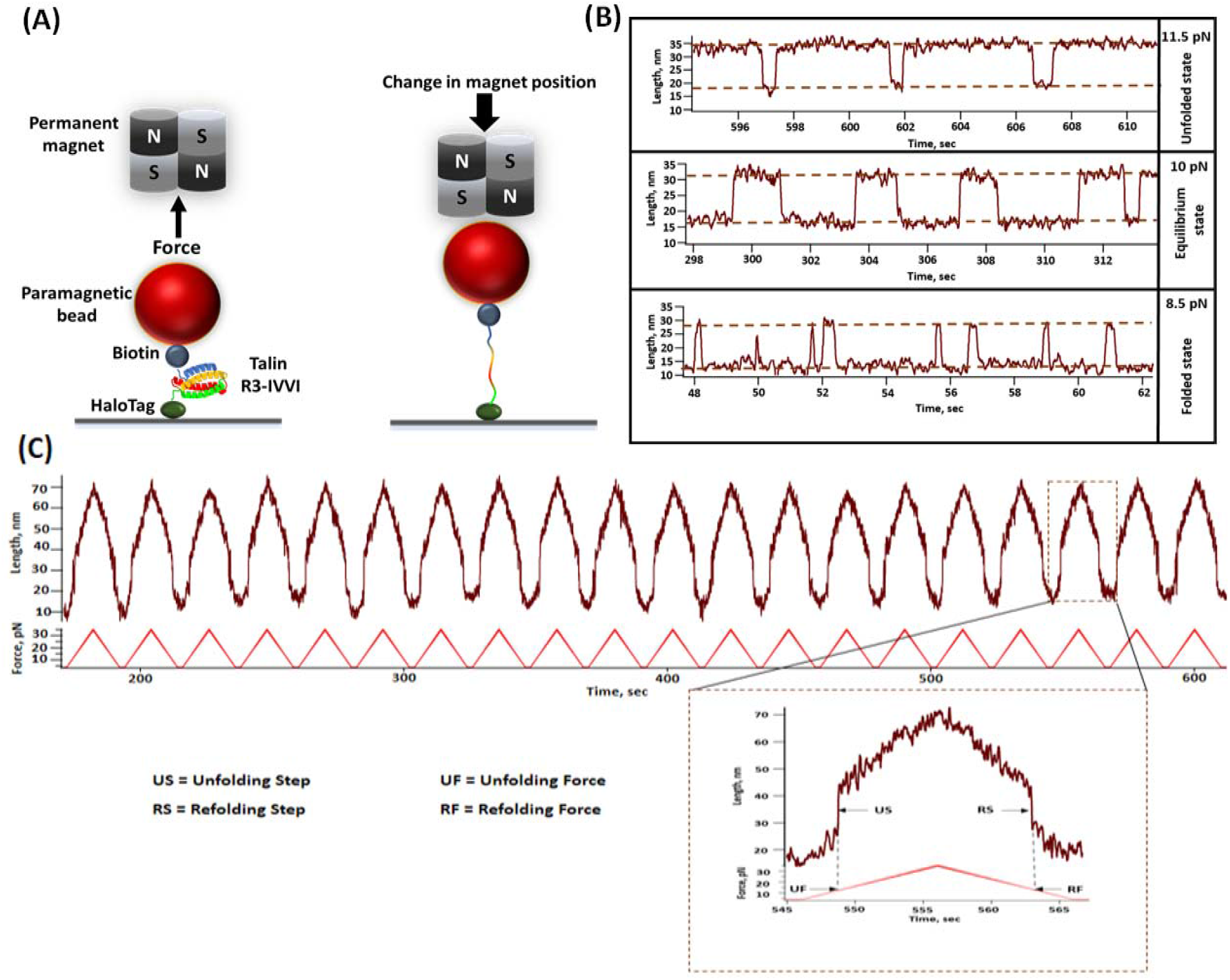
Real time magnetic tweezers set-up to study the folding dynamics and mechanical stability of talin R3-IVVI. **(A) Schematic representation of magnetic tweezers experiment:** The engineered construct of single R3-IVVI domain is flanked between the glass surface and the streptavidin coated paramagnetic bead using halotag chemistry. Applied force is controlled by changing the distance beween paramagnetic bead and permanent magnet. **(B)Force-dependent folding dynamics of talin R3-IVVI domain:** Talin R3-IVVI exhibits a strong force-dependent folding dynamics which is characterized as reversible folding-unfolding transitions, yielding extension changes of ∼ 20 nm. At 11.5 pN force, it is mostly present in unfolded state, while at 8.5 pN, it is mostly in the folded state. At 10 pN, both the states occupy almost equal population. **(C) Force ramp study of talin domain:** Successive force ramp, from 4 to 35 pN at a constant loading rate of 3.1 pN/sec, are applied to observe distinct unfolding and refolding steps at particular forces. In the inset, a single force ramp is magnified where the unfolding step (US) and refolding step (RS) have been extrapolated to the force ramp axis to measure the unfolding force (UF) and refolding force (RF), respectively.

The force clamp technology allowed to provide a distinct force on the substrate and enabled us to observe its hopping between two conformations: folded and unfolded states. Fig. 1B shows three representative trajectories of talin R3-IVVI folding dynamics, and the population of both the folded and unfolded states, at three distinct forces. The folding dynamics was deduced by analysing several of these folding trajectories and dividing the relative population of folded state by the total duration of the observed dynamics as described previously.^20,21^ At 10 pN, the talin R3-IVVI domain displays almost equal proportion of folded and unfolded conformation with an extension change of ∼20 nm; while at 8.5 pN and 11.5 pN, the domain mostly occupies folded and unfolded population, respectively. This force-dependent folding dynamics has been observed to shift in the presence of different chaperones, which is also evident from the change in unfolding and refolding force in force-ramp experiment.

### Chaperones modulate the unfolding and refolding force of Talin

The mechanical stability of R3-IVVI was monitored using the force ramp technology, where we performed the force-ramp experiment by increasing the force from 4 to 35 pN at a constant loading rate of 3.1 pN/s, followed by a successive force-decrease scan with the same loading rate. During each force increase scan, we observed unfolding events as sudden increases in the extension, which is also observed the force-decrease scan as sudden decrease in the extension at particular peak forces. We estimated the peak force by vertically joining the unfolding and refolding steps (US and RS) to the equivalent force in force-ramp axis (as shown in inset Fig. 1C). For the easy understanding, we have also shown a representative force-ramp experiment in the figure below and have described how to estimate unfolding and refolding force (UF and RF). We monitored more than 150 events to quantify the unfolding and refolding force of R3-IVVI domain (Fig. 1C). Interestingly, in the presence of chaperones, the force extension curves are hysteric in nature due to the altered binding dynamics in chaperone-substrate interaction (Supporting Figure 13).

We systematically investigated the chaperone effect on the talin mechanical stability by monitoring their unfolding and refolding force using the force-ramp protocol. The unfolding force of talin has been observed to decrease with the unfoldase chaperone, in compared to the absence of the chaperones (control). For example, the unfolding force is decreased from 10.7±0.2 pN in control, to 7.9± pN in the presence of DnaJ and 7.8±0.3 pN with apo-DnaK (Fig. 2A and 2B). We also performed the experiment with DnaK+DnaJ complex (DnaKJ-ATP) and observed that unfolding force is decreased to 9.4±0.7 pN, implying that DnaKJ complex still works as a holdase under force (Fig. 2C). However, the unfolding force is reverted to that of control in the presence of DnaK+DnaJ+GrpE (DnaKJE-ATP) complex (Fig. 2C). Remarkably, in the presence of 60 µM oxidized DsbA, a known force dependent foldase chaperone,^23^ the mechanical stability of talin increased to 14.7±0.9 pN (Fig. 2D). This chaperone-modulated mechanical stability has also been reconciled from the unfolding/rupture force analysis, where the unfolding forces are plotted against the varying loading rate from 1 to 7 pN/s. We observed that in the absence of any chaperones, the unfolding force at zero loading rate is 7.7±0.01 pN, which decreases to 6.9±0.07 pN in the presence of DnaK while increasing up to 10.4±0.3 pN with DsbA, signifying the altered-mechanical stability of talin with different chaperones (Supplementary Figure 1).

**Figure 2:**
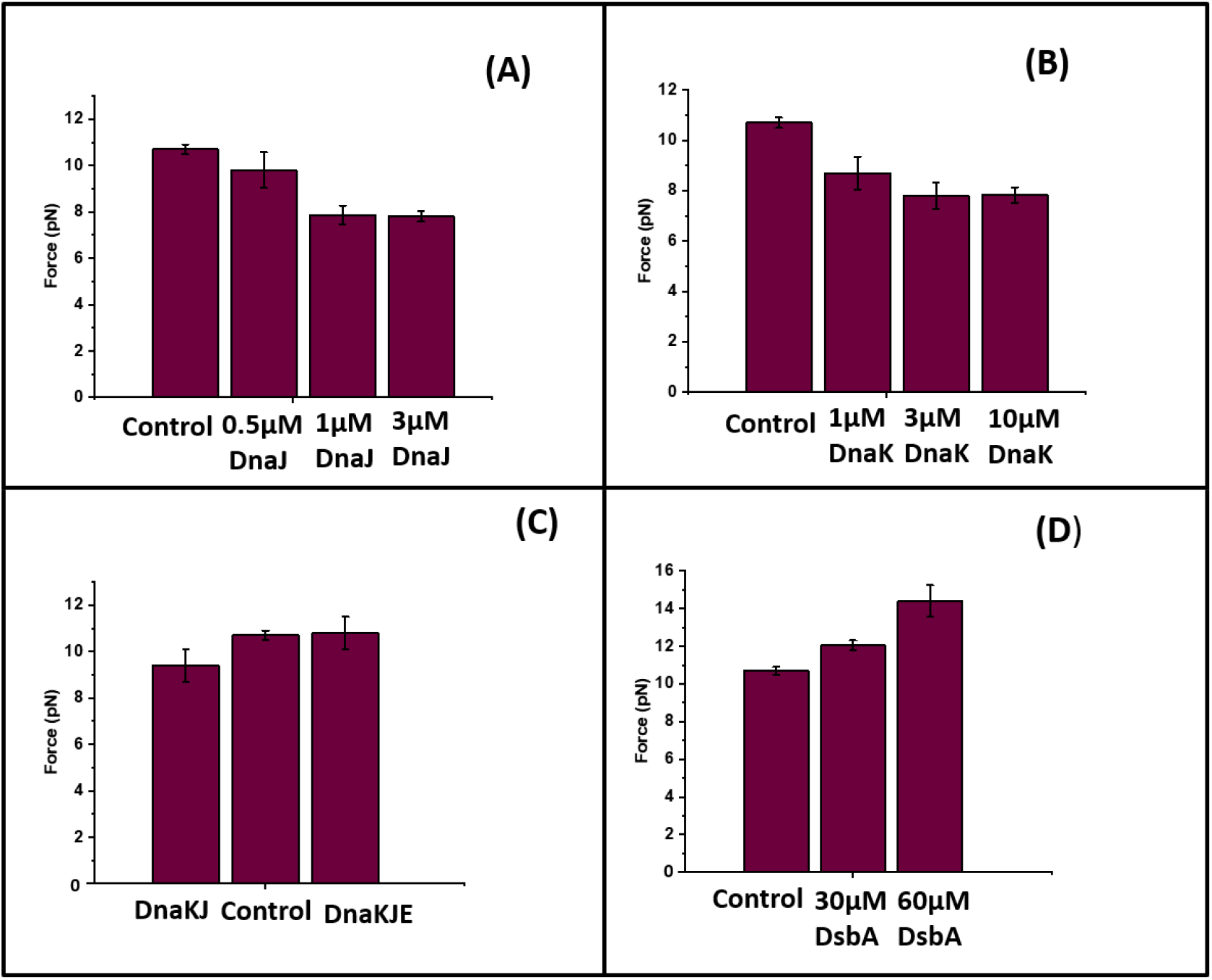
Force ramp experiment detects the chaperone-mediated change in unfolding force of talin: In an unfolding force ramp protocol, the force is increased from 4 to 35 pN at a constant rate of 3.1 pN/sec. **(A) Effect of DnaJ:** Unfolding event in the absence of chaperones (control) is observed at 10.7±0.2 pN, which has been observed to decrease steadily with DnaJ concentration. **(B) Effect of DnaK**: In the presence of DnaK, talin unfolds at lower force and the force decreases with the increasing concentration of DnaK. **(C) Effect of DnaKJ and DnaKJE:** In the presence of DnaK, DnaJ and ATP, talin unfolds at lower force of 9.4±0.7 pN, while upon addition of GrpE (DnaK+ DnaJ+ GrpE+ ATP), it unfolds at a similar force with the control. These experiments have been performed with 10 mM ATP and 10 mM MgCl_2_. The buffer is changed after every 30 minutes with fresh ATP to keep the sufficient supply of ATP. **(D) Effect of DsbA:** DsbA increases the unfolding force in force ramp experiments. Talin domain unfolds at 10.7±0.2 pN force, whereas it increases to 12.1±0.3 pN with 30 µM DsbA and reaches to 14.7±0.9 pN in the presence of 60 µM DsbA. Data points are measured using more than five individual molecules with more than 12 unfolding events. Error bar are represented as s.e.m.

Talin refolding forces, similar to the unfolding forces, has also been observed to alter in the presence of different chaperones. In the absence of any chaperones, the refolding force is 9.9±0.2 pN, which decreases with the unfoldases (Fig. 3A and 3B) and increases in the presence of foldase like DsbA (Fig. 3D). This holdase function is also observed in case of refolding force with DnaKJ-ATP complex, however, addition of GrpE to this complex restores the mechanical stability in talin by increasing the force comparable to that of control (Fig. 3C).

**Figure 3:**
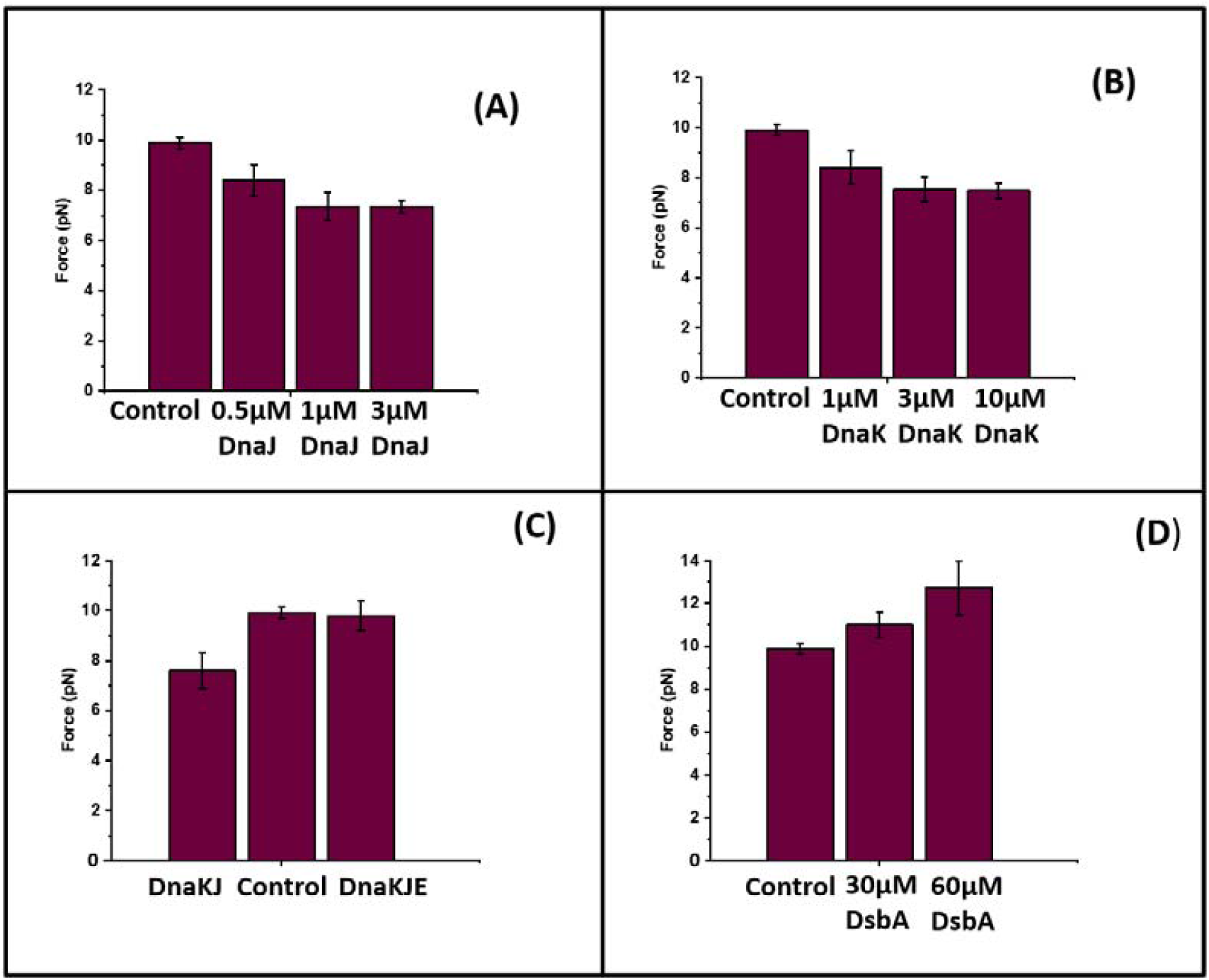
Force ramp protocol detects the chaperon-mediated change in refolding force of talin: In a refolding force ramp protocol, the force is decreased from 35 pN to 4 pN at a rate of 3.1 pN/sec. **(A) Refolding force in the presence of DnaJ:** In the absence of DnaJ, the refolding event of talin is observed at 9.9±0.2 pN and with the gradual addition of DnaJ, the refolding force decreases steadily and becomes saturated at 1 µM DnaJ. **(B) Effect of DnaK:** Similar to DnaJ, DnaK decreases the refolding force of talin domain. **(C) Effect of DnaKJ and DnaKJE on the refolding force of talin:** In the presence of DnaKJ and ATP, R3-IVVI refolds at lower force (7.6±0.7 pN) than control (9.9±0.2 pN), whereas upon addition of GrpE to this mixture (DnaK, DnaJ, GrpE and ATP), it refolds at 9.8±0.6 pN. The buffer is changed after every 30 minutes with fresh ATP to keep the sufficient supply of ATP **(D) Refolding of R3-IVVI in the presence of DsbA:** The refolding force has been observed to increase from 9.9±0.2 pN (control) to 11±0.6 pN in the presence of 30 µM DsbA, and increases further to 12.7±1.2 pN with 60 µM DsbA. Data points are measured using more than five individual molecules with more than 12 refolding events. Error bars are represented as s.e.m.

### Unfolding and refolding kinetics of talin observed under force in presence and absence of chaperones

Dwell time analysis at different force ranges, in the presence of different chaperones, suggest that talin optimally manifests the folding dynamics at a force, where both the folded and unfolded states are equally populated (Supplementary Figure 2 to 6). This reflects to its unfolding and refolding rates during chaperone interactions. We calculated the unfolding and refolding rates of talin, as described previously.^20^ The rates were derived from the inverse of averaged folding and unfolding dwell times (*s*^*-1*^) at particular forces, as determined from the exponential fit to the dwell time distribution. From these measured rate values in mechanical chevron plots, we obtained a linear relationship between their ln values and the applied force, and fitted with a Bell-like equation (eq.2 and 3). Since talin exhibits folding-unfolding transitions within a small force regime of 2 pN, the data could be explained by Bell model.^20^.

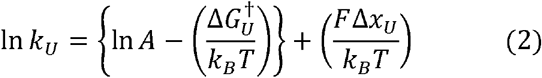

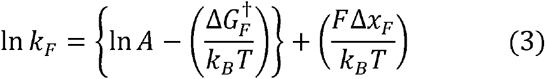

*F* is the applied force, for which the free energy barrier is reduced by *FΔx, Δx* represents the distance to transition state for folding, (*Δx*_*F*_) and unfolding (*Δx*_*U*_). *ΔG*^*†*^_*U*_ and *ΔG*^*†*^_*F*_ are the height of free energy barrier at zero force for unfolding and folding, respectively. *k*_*F*_ and *k*_*U*_ are the folding and unfolding rate constants, *k*_*B*_ Boltzmann constant, *T* is the temperature at kelvin scale and *A* is pre-exponential factor, 10^6^ sec^-1^.^24,25^ The cross-point of unfolding and refolding rates is referred as intersection force, where talin exhibits equal unfolding and refolding rates. However, chaperone interactions perturb the intersection force of talin to different forces such as, in control, the intersection force is 9.7 pN (Fig. 4A), while holdase like DnaJ or apo-DnaK downshift this force to 7.9 pN and 6 pN, respectively (Fig. 4B and 4C). By contrast, the mechanical foldase like DsbA increases it to 14.9 pN (Fig. 4D). Due to the pronounced mechanical effect of chaperones on talin stability, their force ranges are also observed to change significantly, which is evident from the fraction folded analysis (Supplementary Figure 12). We also monitored the intersection force in the presence of chaperone complexes at different nucleotide states (Supplementary Figure 7, 8 and 11).

**Figure 4:**
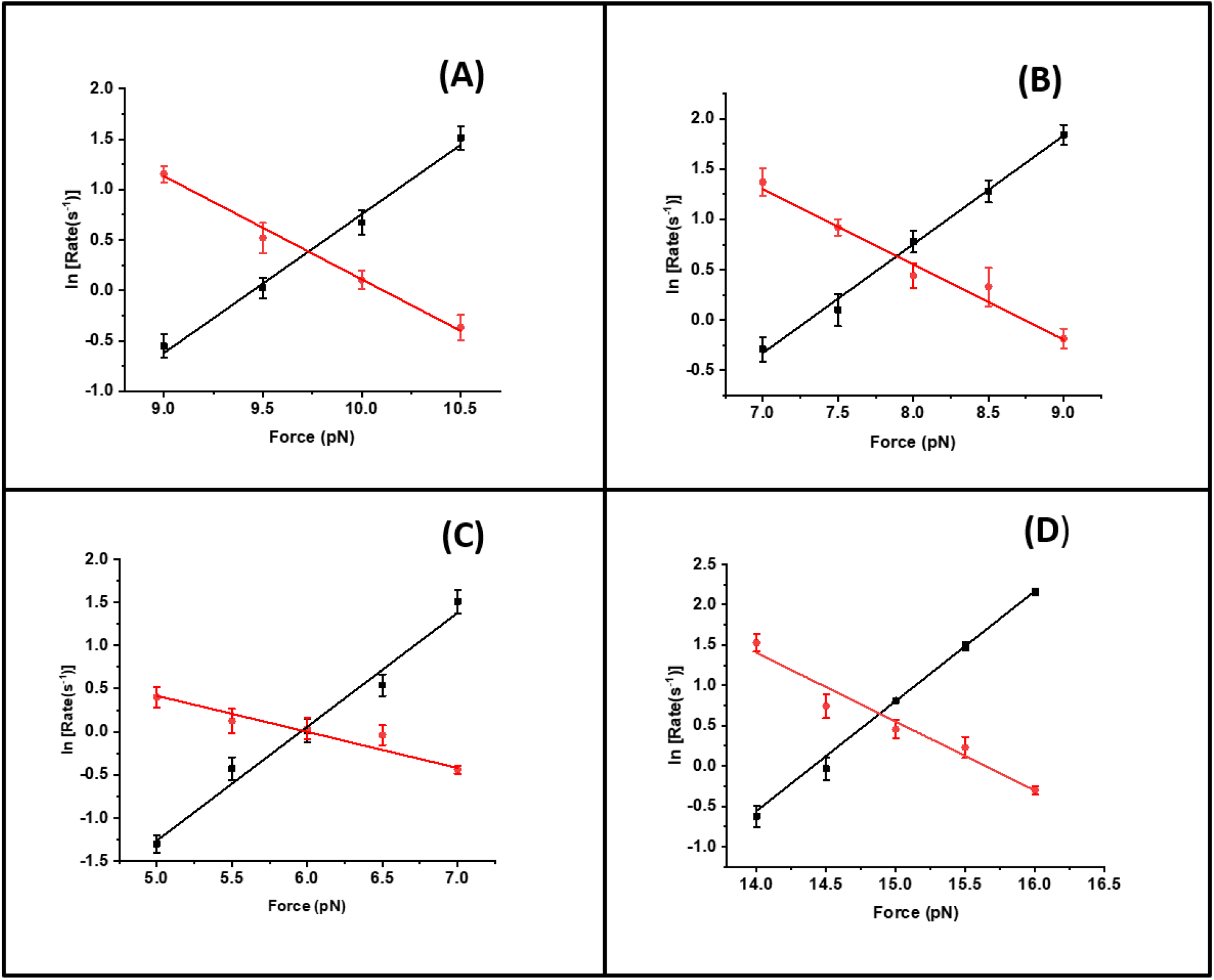
Unfolding and refolding rates of talin domain: **(A) Unfolding and refolding rates in the absence of chaperones (control):** Unfolding (black) and refolding (red) rates are plotted as a function of force. **(B) Effect of DnaJ on unfolding and refolding rates:** Unfolding (black) and refolding (red) rates of talin in the presence of 1 µM DnaJ. Data points are calculated using more than six protein molecules. Error bars are relative error of log. **(C) Effect of DnaK**: Refolding (red) and unfolding (black) rates of R3-IVVI at different forces, in the presence of 3 µM DnaK. Data points are calculated using more than five protein molecules. Error bars are relative error of log **(D) Effect of DsbA:** Unfolding (black) and refolding (red) rates of R3-IVVI are plotted against the force, in the presence of 60 µM DsbA. Data points are calculated using more than four individual molecules per force. Error bars are relative error of log.

Additionally, from the chevron plots analysis, we determined the height of the energy barriers and transition state distance for the unfolded state, allowing us to construct a quantitative chaperone-induced free energy landscape of protein (Table 1). Protein exhibits force-independent unfolding transition due to the rigid folded state, and thus, the unfolding kinetics fits better to Bell-like equation.^26^ However, recent experimental and theoretical studies have suggested that refolding transition is strongly force-dependent due to the compliance variation of the unfolded state and collapse-associated energetics, leading to a non-linear force-dependence on the logarithmic scale, and therefore, Bell model is not a good approximation for refolding kinetics.^27^ This is because unfolded states are highly flexible polypeptide chain (or soft polymer) and thus, are easily subjected to deformation by external force, perturbing its mechanical compliance.^8,27–30^

**Table 1:**
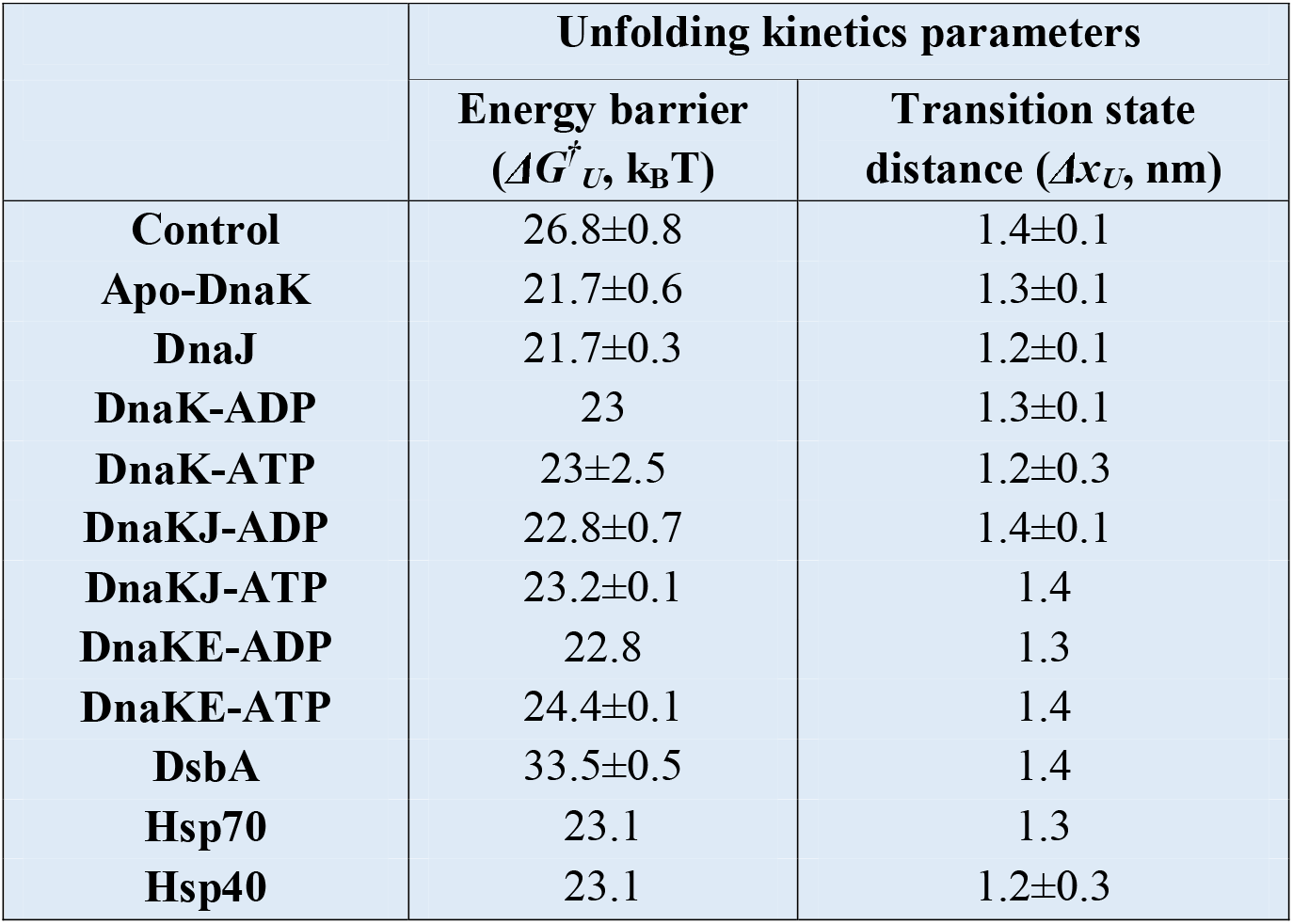
The parameters are obtained by fitting the unfolding rates to Bell model (see Fig 4 and 6). ΔG^†^_U_ represents the height of the free energy barrier and Δx_U_ is the distance from the folded state to the unfolding transition state along the reaction coordinate. Data points are calculated using more than four individual molecules per force. Errors bars are represented as s.e.m.

### Effect of mechanical chaperones on the WT talin domain

To understand the effect of chaperones on wild type (WT) R3 domain, we measured the intersection force of R3-WT in the presence of different chaperones. Due to the low mechanical stability, the intersection force of R3-WT domain in the absence of any chaperones is ∼6 pN. We observed holdase and foldase chaperones can change this force. For instance, in the presence of holdase chaperone such as DnaJ or DnaK, the intersection force decreased to 4 pN, while addition of foldase chaperone DsbA increased it to 10 pN. For the DnaKJE complex, similar to R3-WT, it exhibits intersection force of 6.2 pN (Supplementary Figure 9).

### Effect of Hsp70 and Hsp40 on talin mechanical stability

To generalize our observations with the model DnaK chaperone system, we further tested the mechanical stability of talin with the eukaryotic homologues of DnaK and DnaJ system-Hsp70 and Hsp40.^31^ Since these chaperones reside in eukaryotic cytosol, they most likely to interact with talin and thus, could change the talin folding dynamics by modulating their mechanical stability. Interestingly, we observed Hsp70 and Hsp40, very similar to DnaK and DnaJ, act as mechanical unfoldases and reduce the mechanical stability of talin by changing their unfolding force from 10.7±0.2 pN to 7.7±0.5 and 8.3±0.4 pN, respectively (Fig. 5A) and the unfoldase activity is also evident from the changes in refolding force (Fig. 5B). Additionally, Hsp70 and Hsp40 has been observed to significantly shift the talin folding dynamics to lower force regime (Supplementary Figure 10), which has been reconciled from the intersection force in chevron plot analysis. For example, the intersection force decreases from 9.8 pN to 7.8 pN in the presence of Hsp40 (Fig. 6A) and similarly, 7.2 pN with Hsp70 (Fig. 6B).

**Figure 5:**
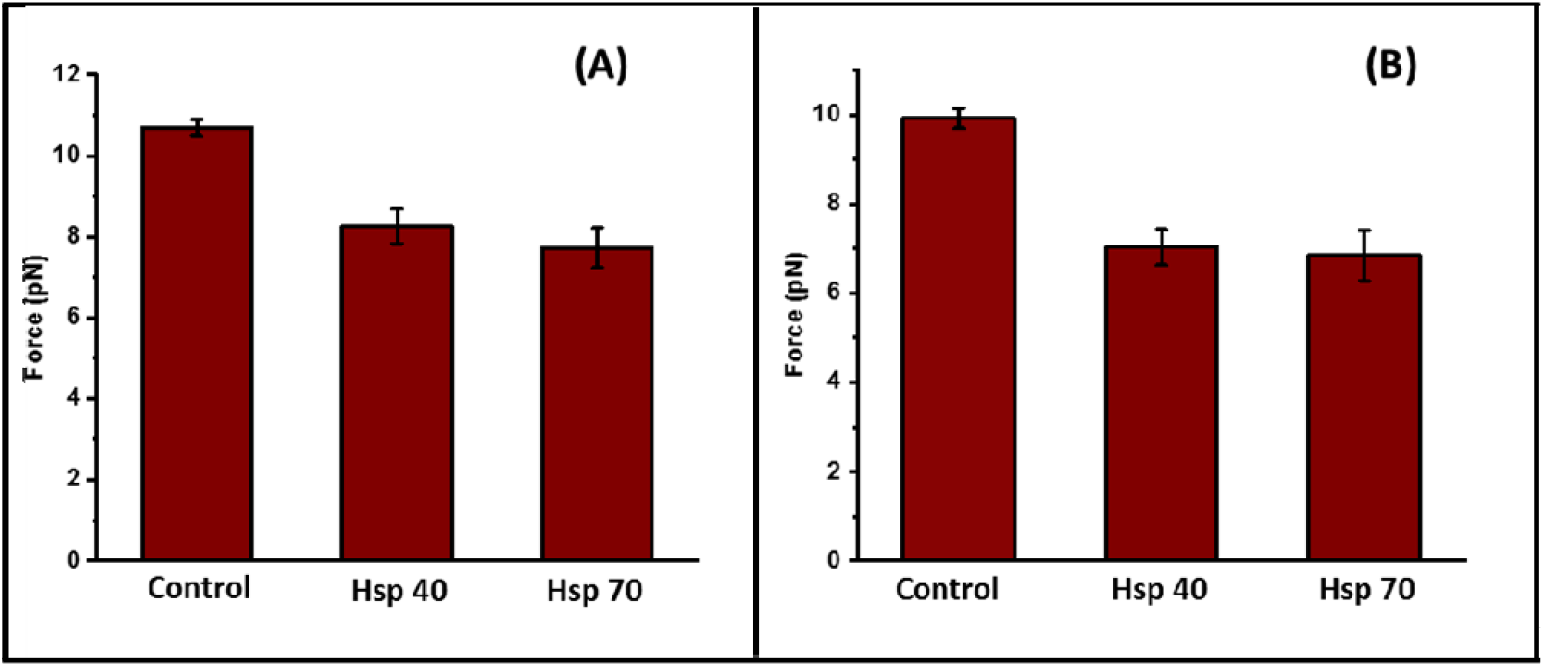
Mechanical stability of talin with Hsp40 and Hsp70: Hsp70 and Hsp40 chaperones decrease he mechanical stability of talin by changing their unfolding and refolding force. **(A) Unfolding force:** In the absence of any chaperones (control), the unfolding force of talin is 10.7±0.2 pN, which has been observed to decrease to 8.3±0.4 pN and 7.7±0.5 pN with Hsp40 and hsp70, respectively. Data points are measured using more than five individual molecules with more than 12 unfolding events. Error bars represent standard error of mean (s.e.m.). **(B) Refolding force:** Similarly, the refolding force has been observed to shift from 9.9±0.2 pN (control) to 7±0.5 and 6.8±0.6 pN with Hsp40 and Hsp70, respectively. Data points are measured using more than five individual molecules with more than ten refolding events. Error bars are represented as s.e.m.

**Figure 6:**
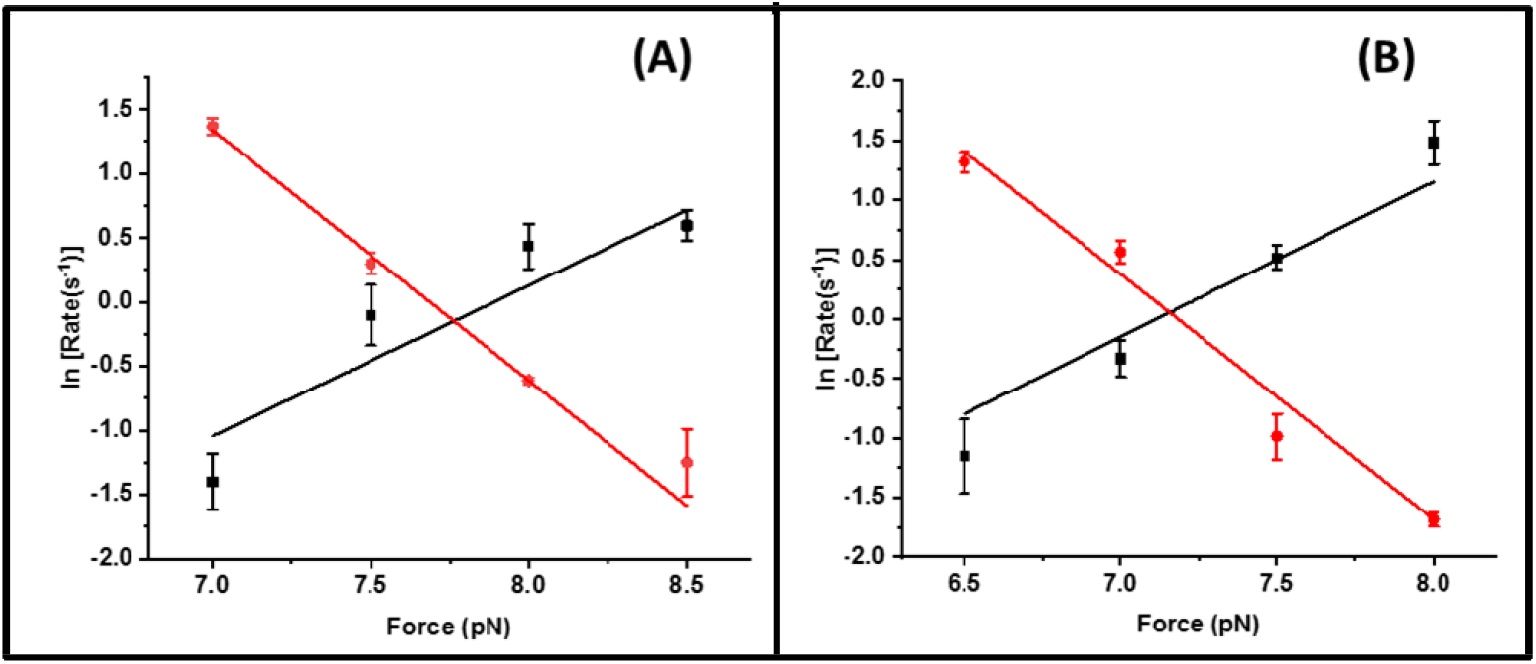
Talin kinetics with Hsp40 and Hsp70: The unfolding and refolding kinetics are plotted as a function of force, in the presence of Hsp40 and Hsp70. Unfolding rate increases and refolding rate decreases with the force and the cross-point of these two rates is defined as intersection force. **(A) Hsp40:** The intersection force in the absence of any chaperones (control) is 9.8 pN, however, has been observed to shift to 7.8 pN with Hsp40. **(B) Hsp70:** Similarly, in the presence of Hsp70, the intersection force of talin is 7.2 pN. This certainly signifies that Hsp40 and Hsp70 decrease the mechanical stability in talin under force. Data points are calculated using more than four individual molecules per force. Error bars are relative error of log.

## Discussion

Force response of talin domains have been well characterized using force spectroscopy and R3 domain has been observed to be the mechanically weakest domain,^8^ exhibiting folding-unfolding dynamics at ∼5 pN. This mechanical instability is originated due to unzipping force geometry and a unique threonine belt at hydrophobic core.^10^ Though R3 domain unfolds at <5 pN force, an energy cost of above 10 k_B_T is required for the unfolding of the domain due to the large unfolding extension of ∼20 nm.^33^ Interestingly, chaperones could modulate the mechanical response of the talin domains, however, the underlying physical mechanism of these interactions under force still remains elusive. Till now, detailed molecular mechanism of diverse protein-chaperone interactions has been illustrated by different groups^34–40^, nevertheless, no studies have elucidated the effect of chaperones in modulating the mechanical stability of force-sensing proteins in FA, which could impact the FA dynamics under physiological force regime.

Magnetic tweezers technology, using fore-clamp protocol, allows us to monitor the chaperone activity on the mechanical stability of talin. We observed that apo-DnaK reduces the mechanical stability of talin domain by decreasing the unfolding force to ∼6 pN and acts as a strong unfoldase. This observation is consistent to the ‘closed’ DnaK conformation in the apo state, which has higher affinity for the peptide substrate, stabilizing it at unfolded state. However, this unfoldase activity is relieved upon the ATP addition, increasing the unfolding force to 7.5 pN. This indicates a lower affinity of DnaK for the peptide substrate in an ‘opened’ conformation during the ATP-bound state.^41^ This ATP-bound DnaK can be cycled between two states: either it remains ATP bound or its ATP hydrolyses into ADP; and this cycle is regulated by two other co-chaperones-DnaJ and GrpE.^31,42–44^ This nucleotide cycle allows DnaK binding to unfolded substrate in both the weaker and tighter binding manners. We observed that addition of DnaJ to DnaK-ATP complex again decreased the unfolding force of substrate protein, reducing its mechanical stability. It is well-known that DnaJ facilitates the ATP hydrolysis^45^, nevertheless, the rationale behind the decreased mechanical stability in the presence of DnaKJ-ATP complex is difficult to ascertain. Interestingly, we monitored the mechanical stability in the presence of only DnaJ and observed that unfolding force is 7.5 pN, as similar to DnaK-ATP complex. Thus, decrease in the mechanical stability is probably caused by the ATP hydrolysis induced by DnaJ. To reconcile this, we also measured the mechanical stability of talin domain in the presence of DnaKJ-ADP and observed that the unfolding force is again decreased to ∼6 pN. Therefore, from these observations, it is evident that ATP binding facilitates the substrate release from the DnaK, resulting in relatively higher unfolding force, while ATP hydrolysis is important for stabilizing the unfolded substrate within the closed DnaK, leading to lower unfolding force. We further added GrpE to both DnaK-ATP and DnaK-ADP complexes and observed the similar pattern in the unfolding force-higher unfolding force in the presence of DnaK, GrpE and ATP and lower force in the presence of GrpE and DnaK-ADP, signifying the accelerated ATPase activity of DnaK in the presence of GrpE,^46^ even higher than in the presence of DnaKJ complex with either ATP or ADP. Finally, upon the addition of the DnaKJE complex with ATP, we observed the unfolding force is same as in the absence of any chaperones, indicating that the complex does not exhibit any additional effect on the unfolding force. These forces are confirmed from the intersection force by mechanical chevron plot analysis, where the unfolding-refolding rates has been plotted against varying forces (Supplementary Figure 7, 8 and 11).

Furthermore, we observed that although chaperones are not able to change the transition state distance significantly, they reshape the free energy landscape by changing the height of the unfolding energy barrier with respect to the unfolded state. The holdase chaperone DnaJ stabilizes the unfolded state by decreasing the unfolding free energy barrier (*ΔG*^*†*^_*U*_) from 26.8±0.8 to 21.7±0.3 k_B_T (decreased by ∼5 k_B_T), inclining the mechanical free energy landscape of R3-IVVI towards the unfolded state (Table 1), which can also be explained from the downshifted folding dynamics. However, DnaKJE complex as a well-known foldase,^47–52^ has only been observed to restore the intrinsic folding ability of talin domain as comparable to the absence of any chaperones. Interestingly, DsbA as a foldase chaperone shifts the free energy landscape towards the folded state by increasing *ΔG*^*†*^_*U*_ to 33.5±0.5 k_B_T (increased by ∼7 k_B_T) (Table 1) which in turn increases the force range of folding dynamics. These Changes in the unfolding barrier height values are in well-agreement with previous single-molecule studies of protein folding dynamics. Brujić et al demonstrated the unfolding dynamics of ubiquitin by single-molecule force spectroscopy and observed that average unfolding barrier height of ubiquitin lies within 5-10 k_B_T range.^53^ More recently, an optical tweezers study revealed that zippering of different domains in SNARE complex met only few k_B_T. For example, the energy barrier of NTD and MD domain zippering were found to be 1 and 2 k_B_T.^54^ Additionally, a single-molecule FRET study shown that free energy barrier of CspTm folding lies within 4-11 k_B_T.^24^ The barrier height in PrP protein has also been observed to be 2±1.2 k_B_T (5±3 kJ/mol) from the unfolded state.^55^ Importantly, these small changes in free energy barrier of ≥ 3 k_B_T could describe a single effective free energy barrier between the folded and unfolded states of protein and thus, signifies the two-state model of protein folding.^56^ This chaperone-mediated reshaping of the free energy landscape apparently affects the mechanical stability of the R3 domain, allowing it to perform as a regulatory mechanical band-pass filter within a narrow force frequency, to convert complex mechanical signals. Therefore, from a broader viewpoint, this could largely impact the force transduction into biochemical signalling by changing the force-dependent folding dynamics.

Mechanotransduction events through the focal adhesions are largely dependent on robust and precise mechanical response of molecular force springs including talin, vinculin etc., which decode the mechanical signals by both the force magnitude and their temporal resolution. Talin rod domains have exquisite ability to interpret these signals by force-driven structural dynamicity and concurrent modulation of complex interactome profiles. These rod domains have graded mechanical stability and unfolds sequentially at increasing tension to control the adhesion dynamics.^8,19^ Among them, R3 domain acts as the initial mechanosensor, initiating talin activation and subsequent signalling pathways. At low force (<5 pN), R3 domain engages to RIAM via folded state, and becomes protected against the mechanical stretching. The tension-induced unfolding at ∼5 pN and concurrent RIAM dissociation is an essential criterion for vinculin binding to cryptic R3 helices (vinculin binding site, VBS). Recently, by pressure-dependent chemical shift, it has also been observed that R3 domain is thermodynamically tuned for interacting with either RIAM or vinculin which undergoes through a stepwise structural transition.^2^ This mutual exclusive interaction of RIAM and vinculin to R3 finely calibrates downstream signalling cascades, reflecting the changes in cell morphology.^57^ RIAM recruitment to talin-integrin mechanical linkage results in filopodial protrusion in nascent adhesion that eventually replaced by vinculin in mature adhesion.^9^ The adhesion reinforcement results from the actomyosin contractility which regulate the FA dynamics by tension-dependent binding constant of antagonistic partners. For example, RIAM, DLC1 binding to folded domains, can stabilize talin by increasing the mechanical force threshold for vinculin binding. R3 stabilization has been reported to affect fibroblast matrix rigidity sensing and YAP (yes associated protein) signalling, while destabilized R3 exhibits conformational dynamics and are more prone to tension-induced unfolding.^58^ This unfolding at relatively low force, and subsequent vinculin binding plausibly activate downstream signalling pathways that decrease traction force generation. In mature adhesion, actomyosin contractility perturbs RIAM binding, abrogating its negative effect on RhoA signalling which enhances the tension at growing adhesion. However, as the adhesion matures and engages more proteins to the complex, the resultant force is decreased on individual talin molecules due to multiple parallel linkage. This force reduction on each mechanical linkage, as force-bearing springs, has been observed by FRET tension sensor and has been reported to be decreased in mature focal adhesion complex than nascent adhesion at cell periphery.^59,60^ This means R3 domain again switch to RIAM binding state and reduces the force transmission through successive domains and thus, regulates focal adhesion dynamics. Interestingly, focal adhesion has also been thought to be controlled by active participation of the molecular chaperones with different adhesion proteins, where they could modulate the folding-unfolding transitions of substrate proteins by changing their mechanical stability and therefore, their interaction with other partners. Our study, from a larger viewpoint, suggest the plausible role of chaperone in modulating the folding dynamics of adhesion proteins under force. This might play a pivotal role in mechanotransduction where force triggers conformational changes of mechanosensing proteins, and thereby, regulating downstream signalling pathways (Fig. 7)

**Figure 7:**
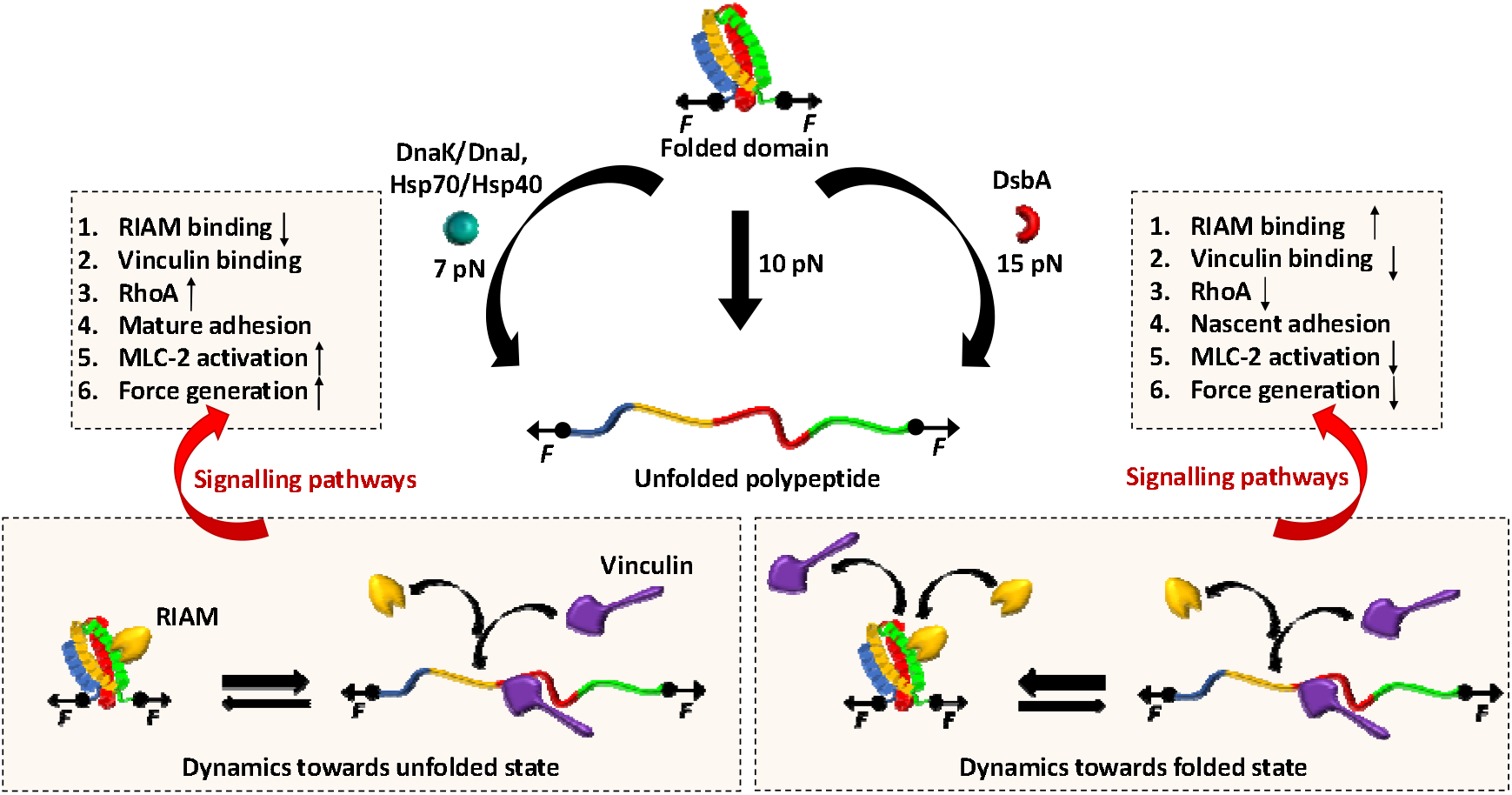
Plausible in vivo effect of Chaperone-modulated mechanical stability of talin: Talin rod domains exhibit strong tension-dependent conformational changes which can further be modulated by both classes of chaperones: unfoldase and foldase. In the absence of chaperones, R3 unfolds at 10 pN force while it shifts to 15 pN or 7 pN, in the presence of foldase and unfoldase chaperones, respectively. This altered mechanical stability could influence cellular physiology by tuning the downstream signaling pathways. Unfoldase such as, Hsp70 and Hsp40 shift the folding dynamics of R3 domain towards the unfolded state, thereby, favouring the RIAM dissociation and vinculin interaction. Subsequently, this induces RhoA signaling, downstream MLC-2 activation and thus, enhances the force generation at growing adhesion to facilitate cell migration. By contrast, foldase (DsbA) shifts the folding dynamics towards folded state which subsequently promotes RIAM binding to the talin domains and localization in nascent adhesion. This negatively regulates the RhoA pathway which in turn abrogates MLC-2 activation, force generation, and the cell migration.

There is emerging evidence to suggest that molecular chaperones could accelerate the folding-unfolding transition of cytoskeletal proteins such as actin and tubulin.^13^ Studies have shown that chaperones like αβ crystallin or calreticulin has played crucial role in deciding mechanical properties of cell, like resilience of cell adhesion, cell spreading, by regulating expression of key focal adhesion proteins.^13,14^ Small heat shock proteins have also been observed to regulate chronic biomechanical stress in heart tissue by regulating dynamics of key FA protein like actin and filamin C.^17^ Additionally, tension-induced ubiquitylation of filamin A promotes chaperone-assisted autophagy which in turn dictate the role of chaperones in focal adhesion assembly.^61,62^ However, the underlying molecular mechanism of these chaperone interactions, during mechanotransduction, has not been reported yet. Interestingly, we observed that Hsp70 and Hsp40, the eukaryotic homologues of DnaK and DnaJ, act as unfoldase by decreasing the unfolding barrier height from 26.8±0.8 k_B_T to 23 k_B_T (Fig. 6), stabilizing its unfolded state. This shifting towards unfolded state is also evident from the lower unfolding force in the presence of Hsp70/Hsp40 chaperones (Fig. 5). During interaction with Hsp70 and Hsp40 in cytosol, talin folding dynamics shifts to lower force range (Supplementary Figure 10) where it is mechanically poised to interact with vinculin or any other unfolded state interactor. Plausibly, in case of higher Hsp70 expressions, talin-vinculin could predominantly interact, perturbing normal force-dependent processes in cells. By contrast, tunnel associated chaperones possess an ability to increase the mechanical stability of their substrates. Such as oxidoreductase DsbA, a well-known model mechanical foldase, has the ability to increase the folding rate and mechanical stability of R3.^23,32,63^ Hence, our results demonstrate that chaperones could modulate the mechanical stability of the FA protein talin, which are characterized by the change in the unfolding free energy barrier. From a broader perspective, our result may have generic mechanistic insight of how talin response, and so their binding kinetics with other interactors, could be affected by chaperone interactions. For example, foldase chaperones facilitate the RIAM binding by shifting the energy barrier towards folded state and slows the stepwise unfolding of the domain, while unfoldase chaperones promote vinculin binding by decreasing the unfolding free energy barrier. This could convert the switch response of R3 domain into a domain-locked response, disrupting the natural downstream signalling pathway to control the diverse cellular processes.

## Materials and Methods

### Expression and purification of talin R3-IVVI

We used the mechanically stable talin R3-IVVI domain as a model substrate, as previously reported in force spectroscopic studies.^20,21^ For purification, the protein construct was transformed into the Escherichia coli BL21 (DE3 Invitrogen) competent cells. Cells were grown in luria broth at 37°C with carbenicillin, till the O.D. becomes 0.6-0.8 at 600nm. The cultures were then induced with 1 mM Isopropyl β-D-thiogalactopyranoside (IPTG, Sigma-Aldrich) overnight at 25°C. The cells were pelleted and re-suspended in 50 mM sodium phosphate buffer pH 7.4, containing 300 mM NaCl and 10% glycerol. Phenylmethylsulfonyl (PMSF) was used as protease inhibitor followed by lysozyme for membrane lysis. After incubating the solution for 20 minutes at 4°C, dissolved pellet was treated with Triton-X 100 (Sigma Aldrich), DNase, RNase (Invitrogen) and 10 mM MgCl_2_ and kept at 4°C in rocking platform. The cells were disrupted in French press and cell lysate was centrifuged at 11000 rpm for 1h. The protein was purified from the lysate using Ni^2+^-NTA column of [KTA Pure (GE healthcare). For in vitro biotinylation of the polyprotein, Avidity biotinylation kit was used and the biotinylated polypeptide was purified by Superdex-200 increase 10/300 GL gel filtration column in presence of Na-P buffer with 150 mM NaCl.^63^

### Expression and purification of Chaperones (DnaJ, DnaK, GrpE and DsbA)

For purification of DnaJ and GrpE protein purification, these constructs were transformed in Escherichia Coli (DE3) cells and grown in luria broth media at 37°C with respective antibiotics (carbenicillin and ampicillin, 50µg/ml). The cells were induced overnight with Isopropyl β-D-thiogalactopyranoside (IPTG, Sigma-Aldrich) at 25°C. Cells were centrifuged and the pellet was re-suspended in 50 mM sodium phosphate buffer, 10% glycerol and 300 mM NaCl, pH 7.4. The solution was then incubated with PMSF and lysozyme. DNase, RNase, MgCl_2_ and Triton-X 100 was mixed subsequently at 4°C. Then the cells were disrupted in French press and the lysate was extracted and purified using Ni^2+^-NTA affinity column of [KTA Pure (GE healthcare). We used 20 mM imidazole containing buffer as binding buffer and 250 mM imidazole containing sodium phosphate buffer as elution buffer. We purified full-length DnaK chaperone from the cell lysate using the Ni^2+^-NTA column, pre-loaded with Mge1.^64^ Then the DnaK protein was eluted with buffer containing 2 mM ATP.^64^ For DsbA, the pellet was dissolved in 50 mM Tris buffer with 1 mM EDTA and 20% (w/v) sucrose, pH 8.0. After 20 minutes of ice incubation the mixture was provided with DNase, RNase and PMSF. Then centrifuged the solution followed by collecting both the supernatant (S-1) and pellet. The pellet was dissolved in 20 mM Tris solution and again centrifuged and the supernatant (S-2) was collected. The purification was done by purifying these two supernatants with 1 M NaCl containing Tris buffer using Hi-Trap Q-FF anion exchange column of ÄKTA Pure (GE healthcare). To maintain the oxidation activity of DsbA 0.3% H_2_O_2_ was added to the purified protein for overnight. The protein was further purified by size exclusion chromatography using Superdex-200 increase 10/300 GL gel filtration column in presence of 150 mM NaCl.^23^

### Preparation of glass slide and coverslips

During the magnetic tweezers experiment, the glass slides were washed with Hellmanex III (1.5%) solution (Sigma Aldrich) followed by washing with double distilled water. It was then soaked in a mixture containing concentrated hydrochloric acid (HCl) and methanol (CH_3_OH). The slides were then treated with concentrated sulphuric acid (H_2_SO_4_) followed by washing in double distilled water. The glass slides were put into gently boiling water and dried. To activate the glass surface, the glass slides were dissolved in the ethanol solution of 1% (3-Aminopropyl) trimethoxysilane (Sigma Aldrich, 281778) for 15 minutes. Then the glasses were washed with ethanol for removing the unreacted silane and baked them in 65°C. The coverslips were washed with Hellmanex III (1.5%) solution for 15 minutes followed by ethanol and dried in the oven for 10 minutes. After sandwiching the glass and coverslips, the chamber was filled with glutaraldehyde (Sigma Aldrich) for an hour. Then the chambers were flushed with reference beads (2.5-2.9 µm, Spherotech, AP-25-10) and Halo-Tag (O4) ligand (Promega, P6741). To avoid non-specific interaction the glass chambers were washed with blocking buffer (20 mM Tris-HCl, 150 mM NaCl, 2 mM MgCl_2_, 0.03% NaN_3_, 1% BSA, pH 7.4) for 5 hours at room temperature.^18,65^

### Real time magnetic tweezers experiment

The real time magnetic tweezers set up was built on an inverted microscope using 63x oil-immersion objective that is attached with a nanofocusing piezo actuator. A linear voice-coil is located above the sample chamber to control the position of the magnet. Images were acquired using ximea camera (MQ013MG-ON). Details information regarding the bead tracking, image processing, and force calibration are described previously.^22^ We have also provided in detail description of force calibration method in the supplementary information (Supplementary Figure 14). Real time magnetic tweezers experiment was performed with 1-10 nM of protein in a buffer containing 1X PBS buffer (pH 7.2) and 10 mM ascorbic acid.^66,67^ Ascorbic acid was used as antioxidant for preventing the oxidative damage of the protein. After passing the protein through the chamber, streptavidin coated paramagnetic beads (Dynabeads M-270, cat. No. 65305) were passed through the chamber, where they gets adhered to the biotinylated Avi-Tagged protein. Folding and unfolding dynamics of the protein was observed by applying different force ramp and force clamp protocols. For experiments with chaperones, we used 0.5 µM, 1 µM and 3 µM of DnaJ, whereas DnaK was introduced at 1.5 µM, 3 µM and 10 µM concentration. With DnaKJ and DnaKJE complex, we used 1 µM DnaJ and 3 µM DnaK with 5 µM GrpE. During experiments with DnaK, DnaKJ and DnaKJE the buffers were supplemented with 10 mM ATP and 10 mM MgCl_2_. The buffers were exchanged after every 30 minutes with fresh ATP to provide sufficient ATP supply.

## Acknowledgement

We thank Ashoka University for support and funding. S.H. thanks DBT Ramalingaswami Fellowship and DST SERB Core Research Grant for funding. We thank Dr. Koyeli Mapa of Shiv Nadar University for kindly sharing with us the clones of DnaK, DnaJ, GrpE proteins. We thank Prof LS Shashidhara (Ashoka University), and Dr Edward C. Eckels (Columbia University Medical Center, New York) for the critical analysis and discussion of this work.

## Conflict of Interest

The authors declare no conflict of interest.

## Supplementary Information

**Supplementary Figure 1:**
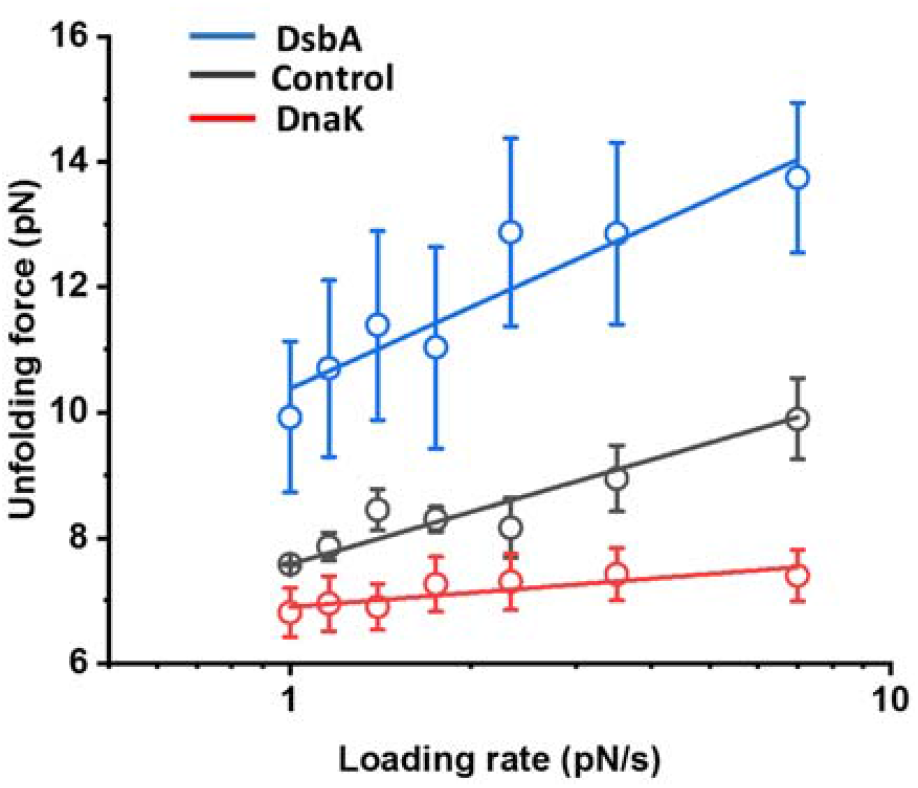
Rupture/unfolding forces of talin at varying loading rates: The unfolding forces are plotted as a function of varying loading rates, ranging from 1 to 7 pN/s. The unfolding forces at zero loading rate has been observed to change in the presence of different chaperones, signifying the change in mechanical stabilities of talin with different chaperones. For example, in the absence of any chaperones, the unfolding force is 7.6±0.1 pN, which has been observed to shift to 6.9±0.1 pN with DnaK and increases to 10.4±0.3 pN. This signifies the increased mechanical stability of talin upon DsbA interaction and lower stability with DnaK unfoldase chaperone. Data points are measured using more than five individual molecules with more than ten unfolding events. Error bars are standard deviation (s.d).

**Supplementary Figure 2:**
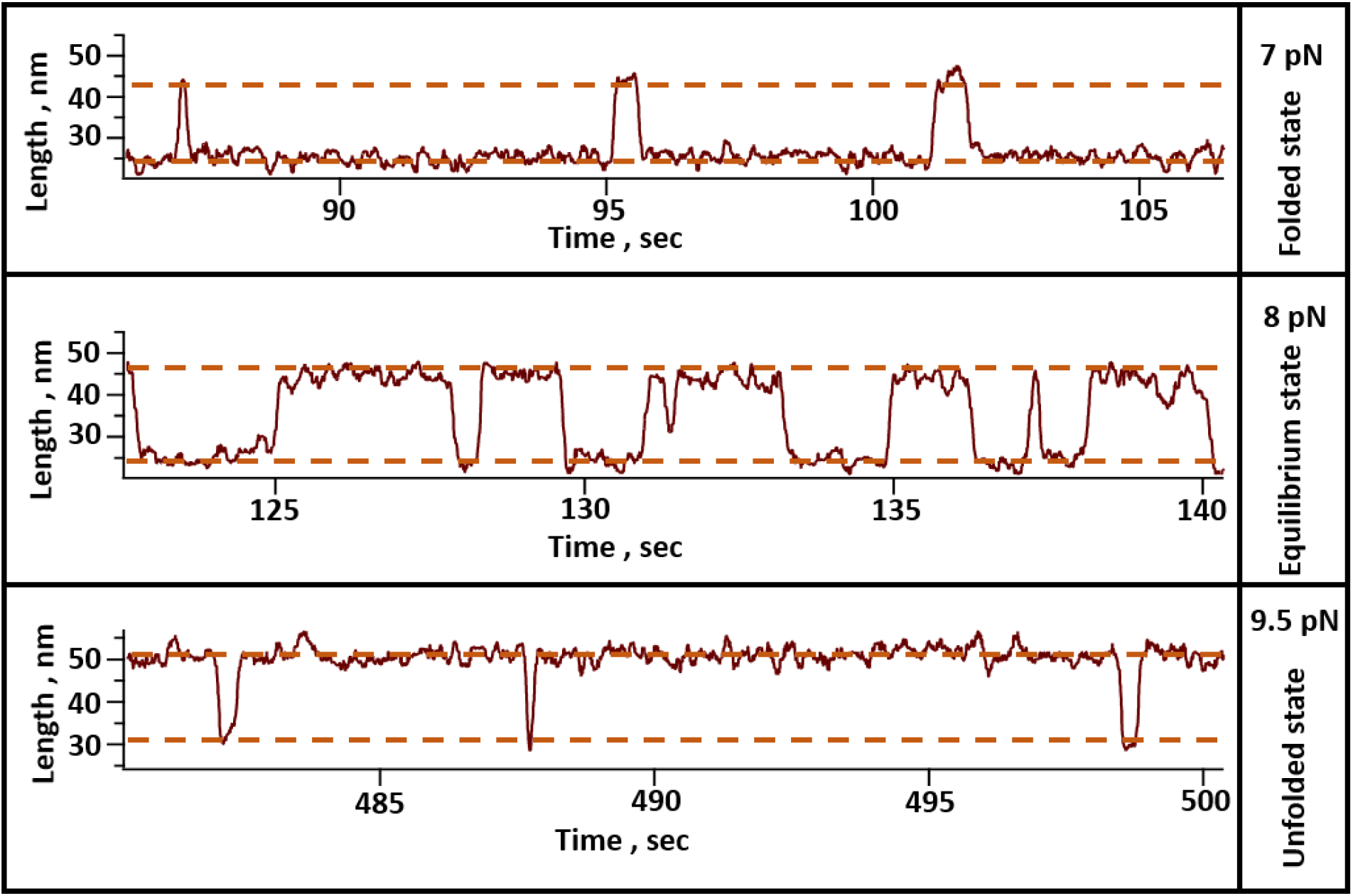
Representative trace of talin R3-IVVI in the presence of 1 µM DnaJ: In the presence of 1 µM DnaJ, the folding dynamics of talin domain shifts towards the lower force regime. At 7 pN, talin mostly stays in the folded state (top trace) and at 9.5 pN force, it mostly stays in the unfolded state. At 8 pN, the domain occupies both the folded and unfolded states.

**Supplementary Figure 3:**
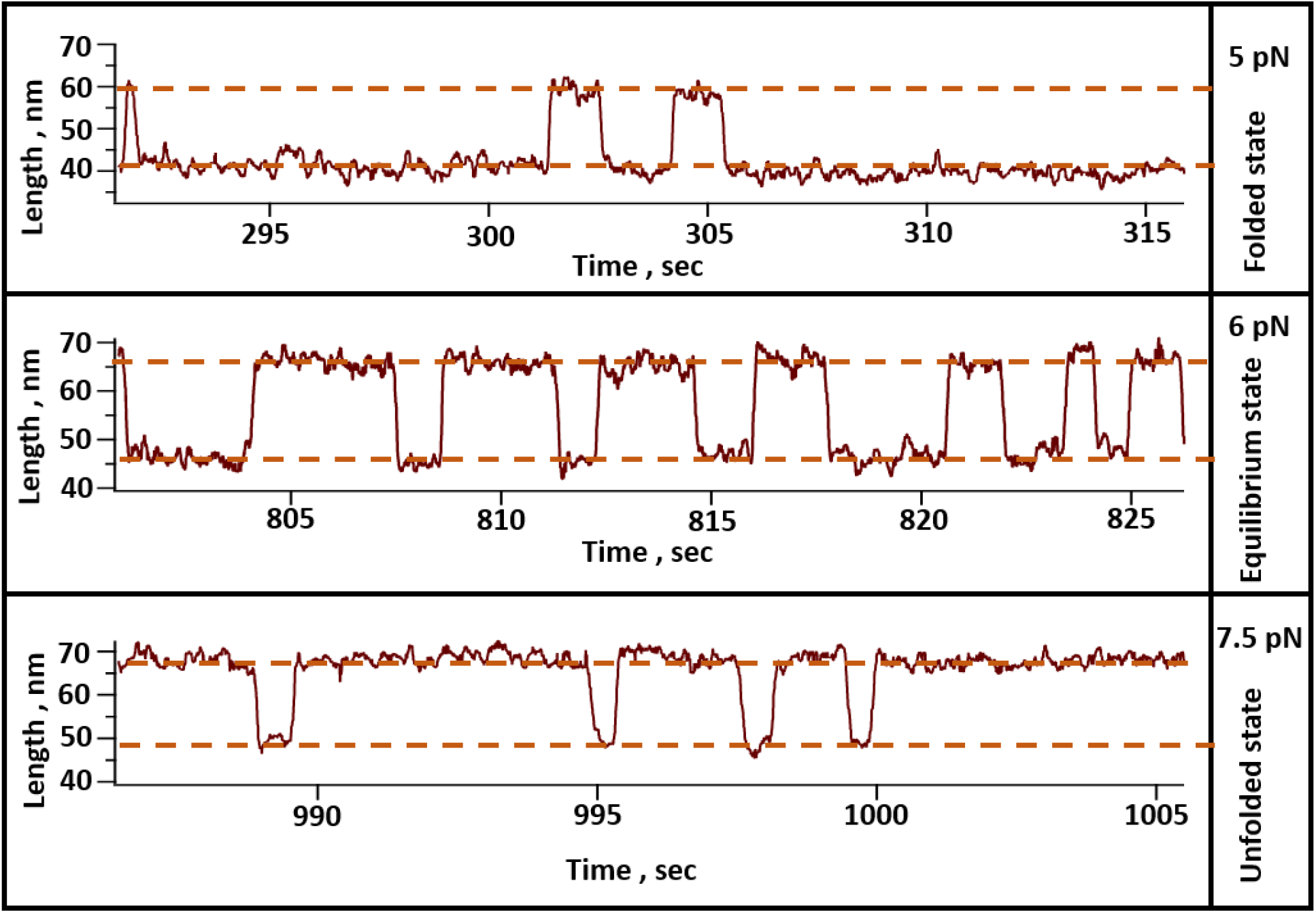
Representative trace of talin in presence of 3 µM apo-DnaK: In the presence of 3 µM DnaK without any nucleotide, the folding dynamics of talin shifts towards the lower force regime. The domain mostly stays in the folded state at 5 pN (top trace), whereas, at 7.5 pN it mostly populates in the unfolded state (bottom trace). At 6 pN force, both the folded and unfolded states are almost equally populated in the domain.

**Supplementary Figure 4:**
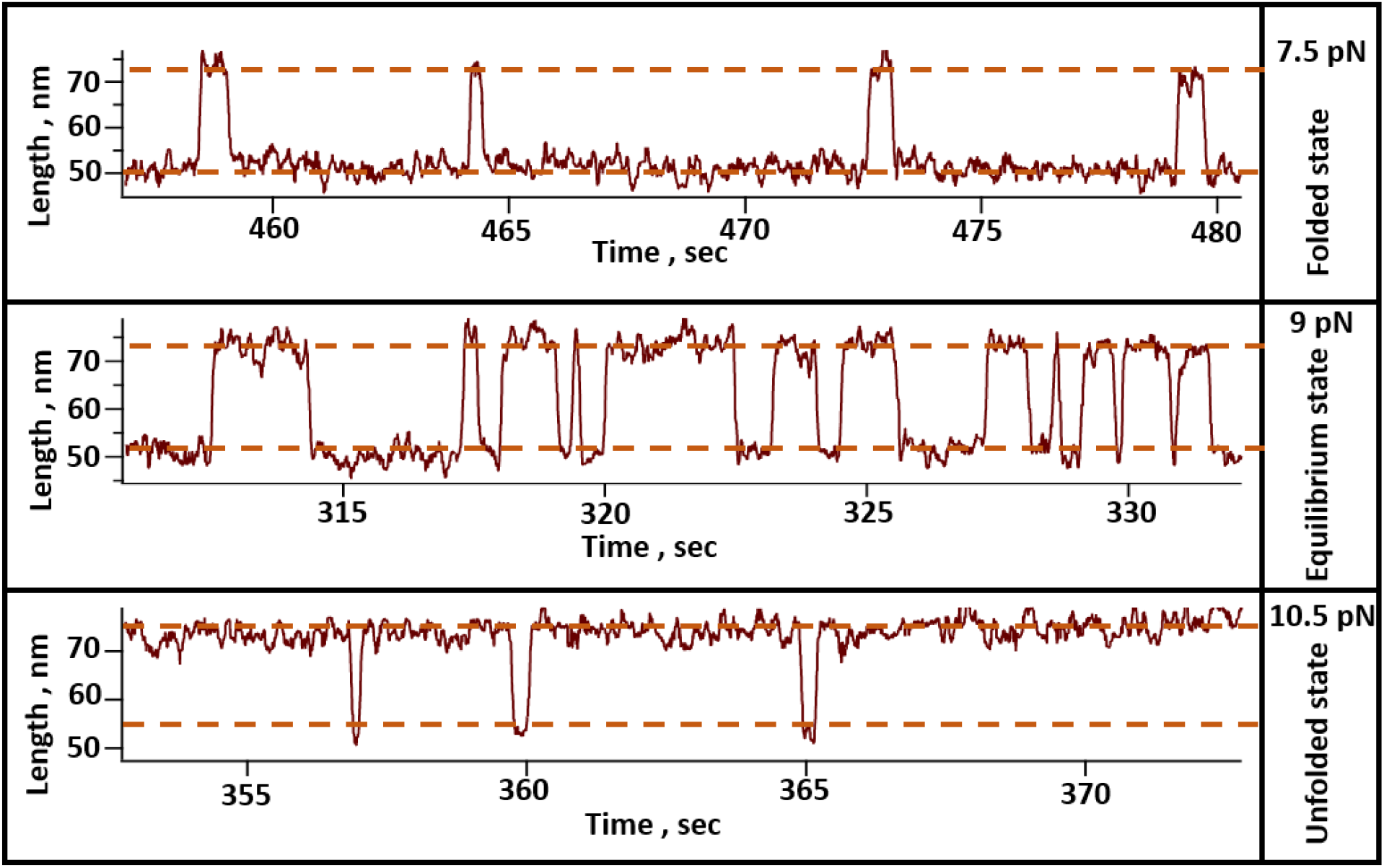
Representative trace of talin in presence of DnaKJ-ATP complex: We monitored the folding dynamics of talin R3-IVVI in the presence of 1 µM DnaJ, 3 µM DnaK and 10 mM ATP and 10 mM MgCl_2_. The buffer is exchanged after every 30 minutes with fresh ATP to keep the sufficient supply of ATP. In this condition, talin mostly populates in the folded state at 7.5 pN (top trace) and mostly in the unfolded state at 10.5 pN (bottom trace). At 9 pN, both the folded and unfolded states are equally populated.

**Supplementary Figure 5:**
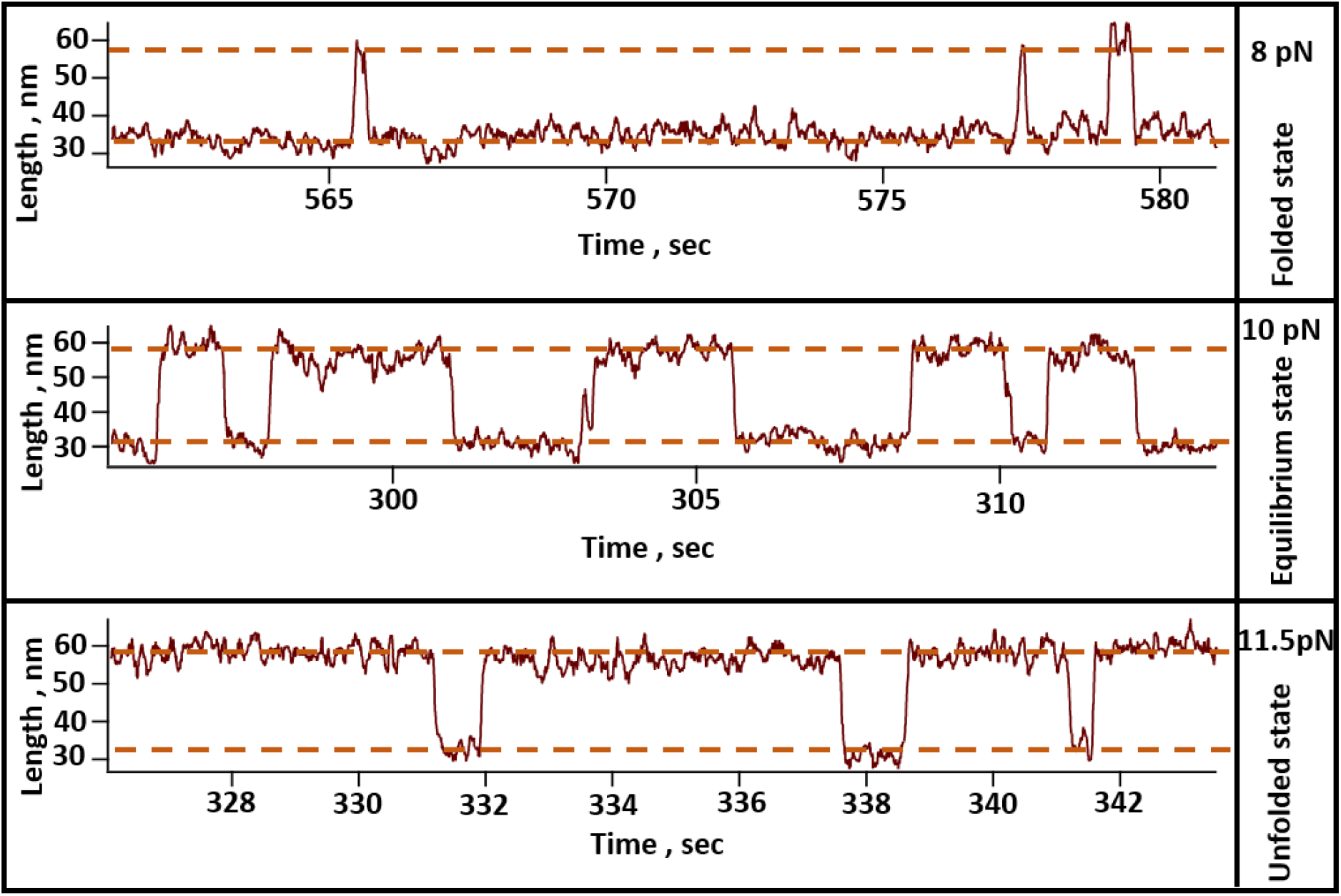
Representative trace of talin domain in the presence of DnaKJE-ATP complex: The dynamics of talin has been monitored in the presence of 1µM DnaJ, 3µM DnaK, 5 µM GrpE and 10 mM ATP and 10 mM MgCl_2_.The buffer is exchanged after every 30 minutes with fresh ATP to maintain the sufficient supply of ATP. DnaKJE complex restores the ability of folding dynamics in R3-IVVI and this domain mostly stays in the folded state at 8 pN (top trace) and in unfolded at 11.5 pN (bottom trace).

**Supplementary Figure 6:**
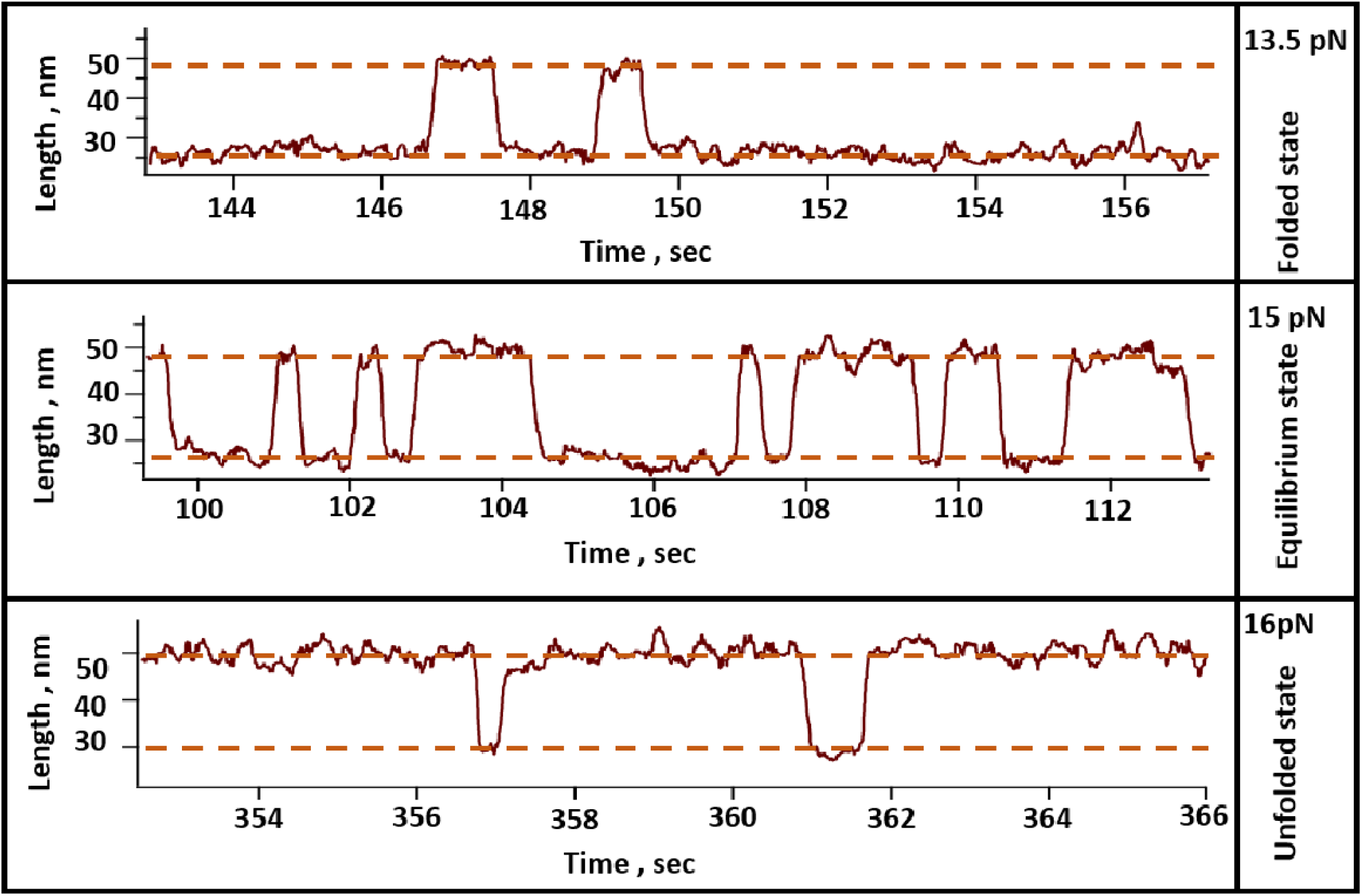
Representative trace of talin R3-IVVI in presence of 60 µM DsbA: In the presence of 60 µM DsbA, the folding dynamics of talin R3-IVVI domain shifts towards the higher force regime. The domain mostly stays in the folded state at 13.5 pN force (top trace), whereas at 16 pN force, it mostly populates in the unfolded state (bottom trace). At 15 pN force, both the folded and unfolded states are equally populated.

**Supplementary Figure 7:**
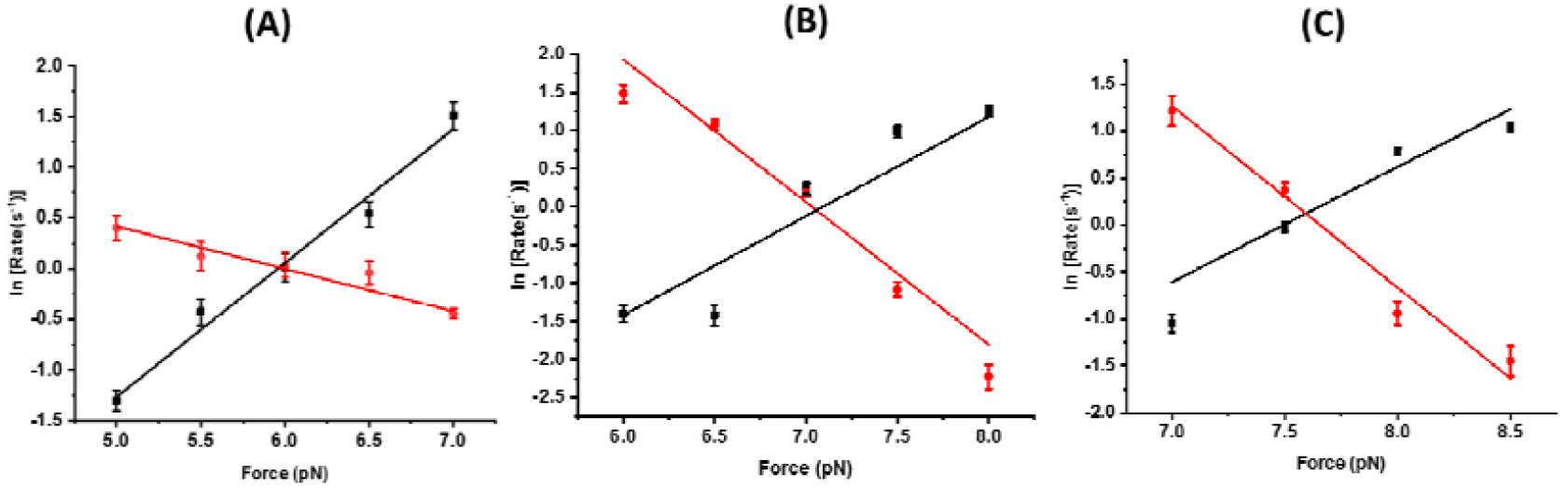
Chevron plots with the different nucleotide states of DnaK chaperone: **(A) Apo-DnaK:** In nucleotide free state or in apo-state of DnaK, the intersection force of talin is 6 pN. **(B) DnaK-ADP:** with DnaK-ADP complex, the intersection force of talin is 7.1 pN. **(C) DnaK-ATP:** in DnaK-ATP state, the intersection force of talin is 7.6 pN. The errors are relative error of log.

**Supplementary Figure 8:**
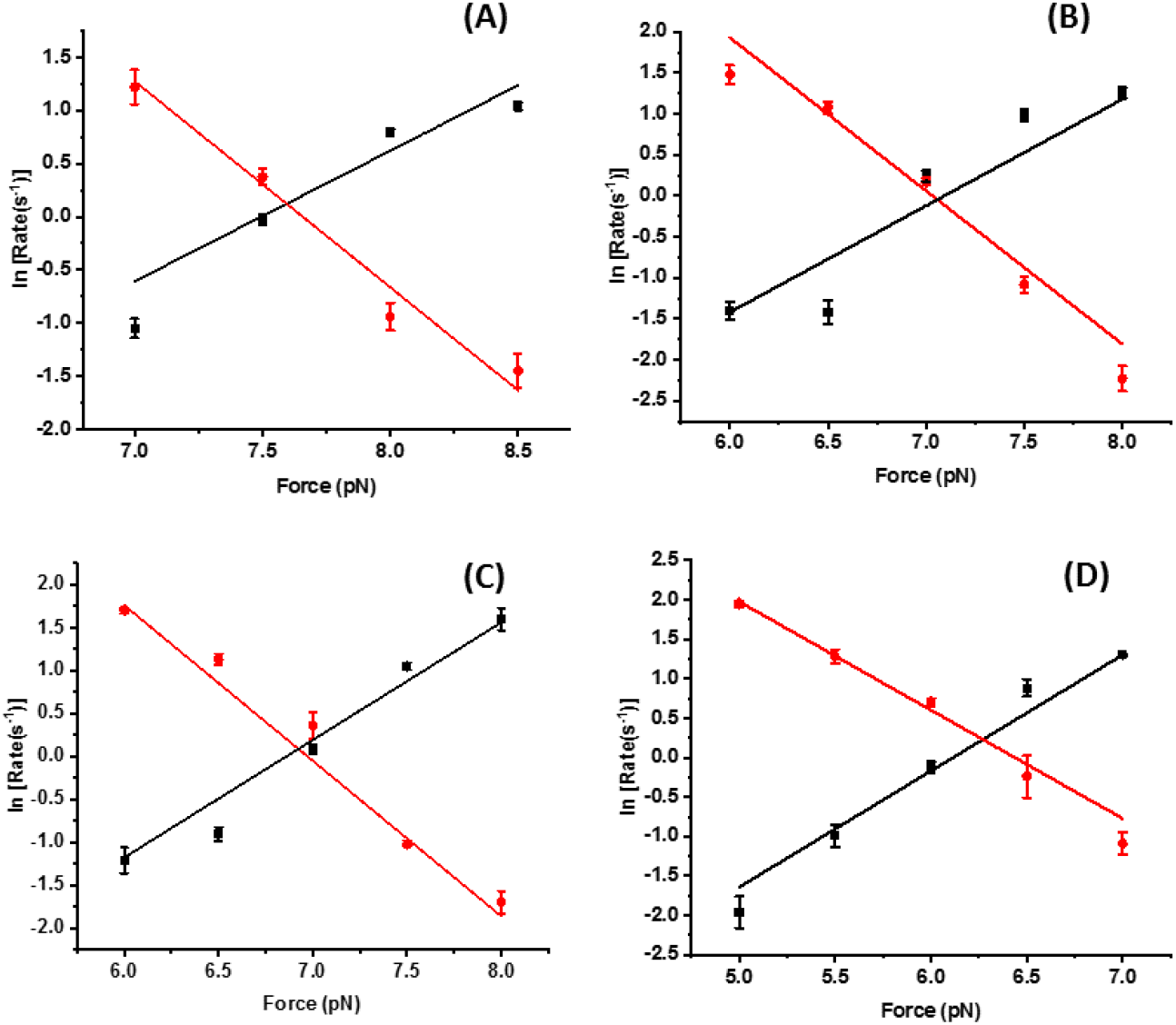
Unfolding and refolding rates with different chaperones: The unfolding and refolding kinetics are plotted as a function of force, in the presence different nucleotide states of DnaK-GrpE. Unfolding rate increases and refolding rate decreases with the force and the cross-point of these two rates is defined as intersection force. **(A) DnaK +ATP:** In the presence of DnaK+ATP, the intersection force is 7.6 pN. Error bars are relative error of log. **(B) DnaK +ADP:** With DnaK+ADP, the intersection force has been observed to decrease from 9.8 pN to 7.1 pN. Error bars are relative error of log. **(C) DnaKJ+ATP:** Similarly, with DnaKJ+ATP complex, the intersection force of talin is 6.9 pN. Error bars are relative error of log. **(D) DnaKJ+ADP:** The intersection force decreases further to 6.2 pN with DnaKJ+ADP complex. Data points are calculated using more than four individual molecules per force. Error bars are relative error of log.

**Supplementary Figure 9:**
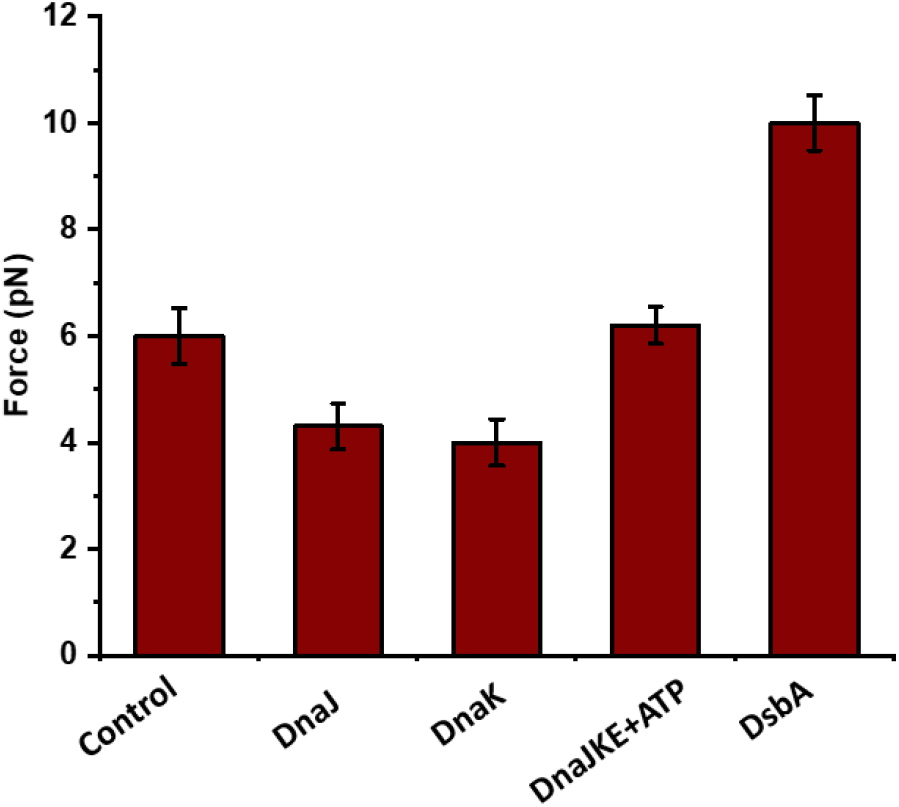
Intersection force of talin WT-R3: Intersection-point forces of talin WT-R3 are measured both in the absence of any chaperones (control) and in the presence of the different chaperones. Intersection-point force is lowered in case of unfoldase chaperones such as, DnaJ and DnaK, while increases with the foldase such as, DsbA. Data points are calculated using more than three protein molecules. Error bars are represented as s.e.m.

**Supplementary Figure 10:**
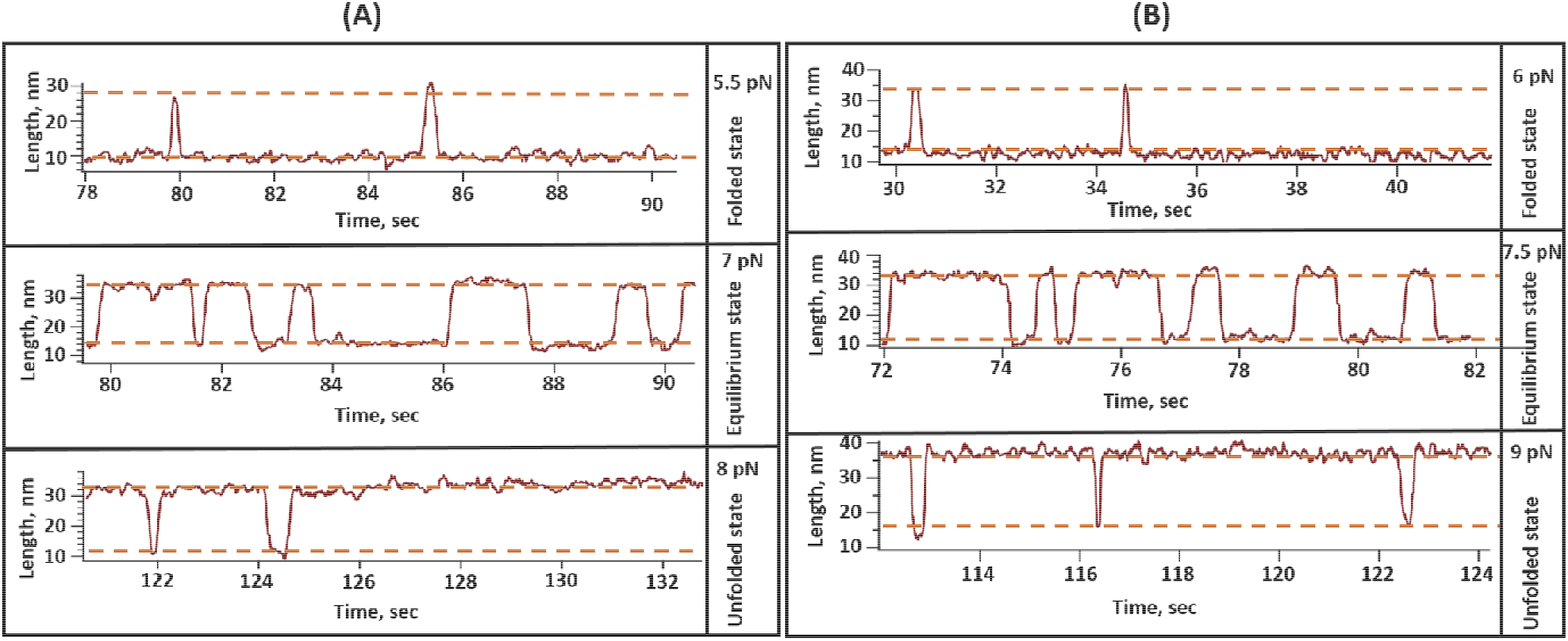
Talin folding dynamics in presence of Hsp70 and Hsp40: **(A) Hsp70:** In the presence of 3 µM Hsp70, talin folding dynamics has been observed to downshift to 7 pN force, with mostly unfolded state at 8 pN and folded state at 5.5 pN. **(B) Hsp40:** Similar to Hsp70, talin equilibrium state shifts to 7.5 pN force with 1 µM Hsp40 and it reaches mostly unfolded state 9 pN and attains mostly folded state at 6 pN.

**Supplementary Figure 11:**
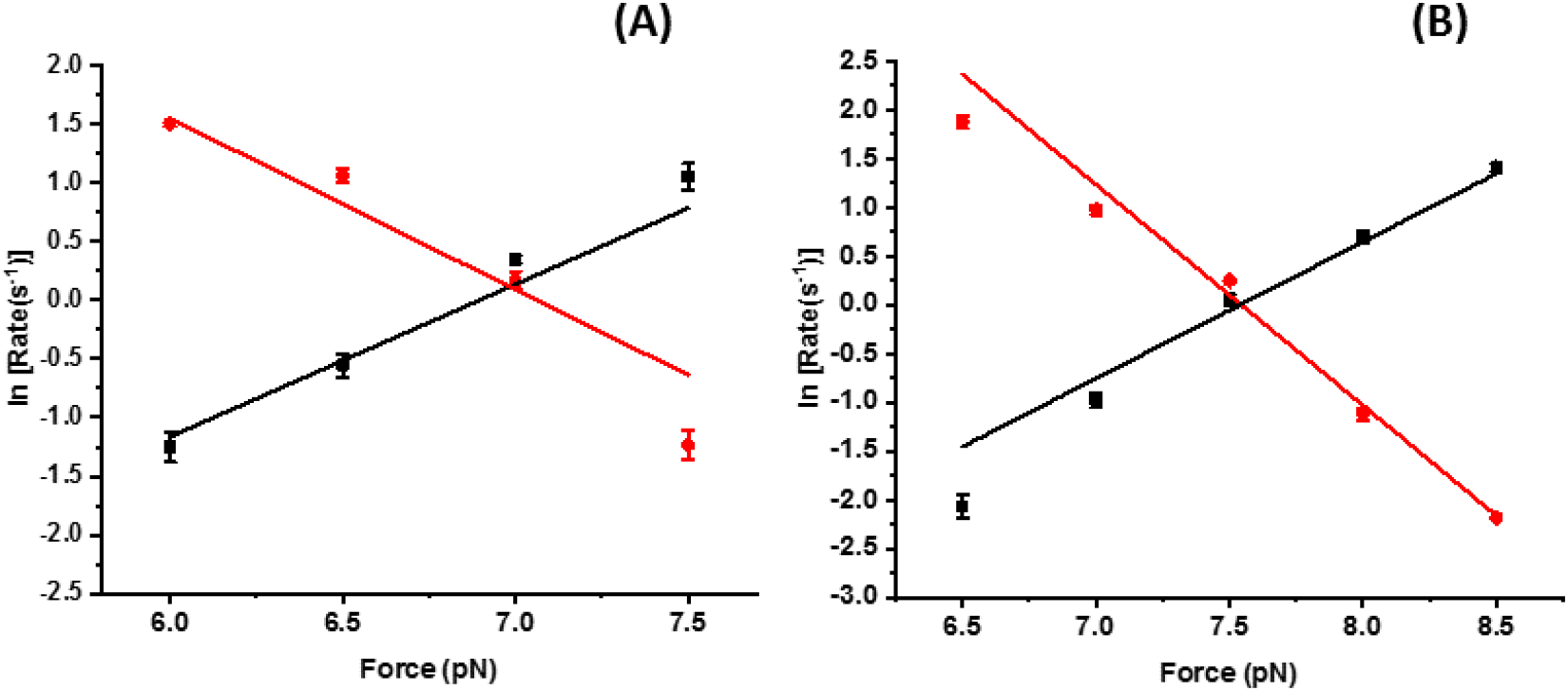
Unfolding and refolding rates with DnaK+GrpE at different nucleotide states: **(A) KE-ADP:** the intersection force is 7 pN. **(B)KE-ATP:** the intersection force is 7.5 pN. Data points are calculated using more than four individual molecules per force. Error bars are relative error of log.

**Supplementary Figure 12:**
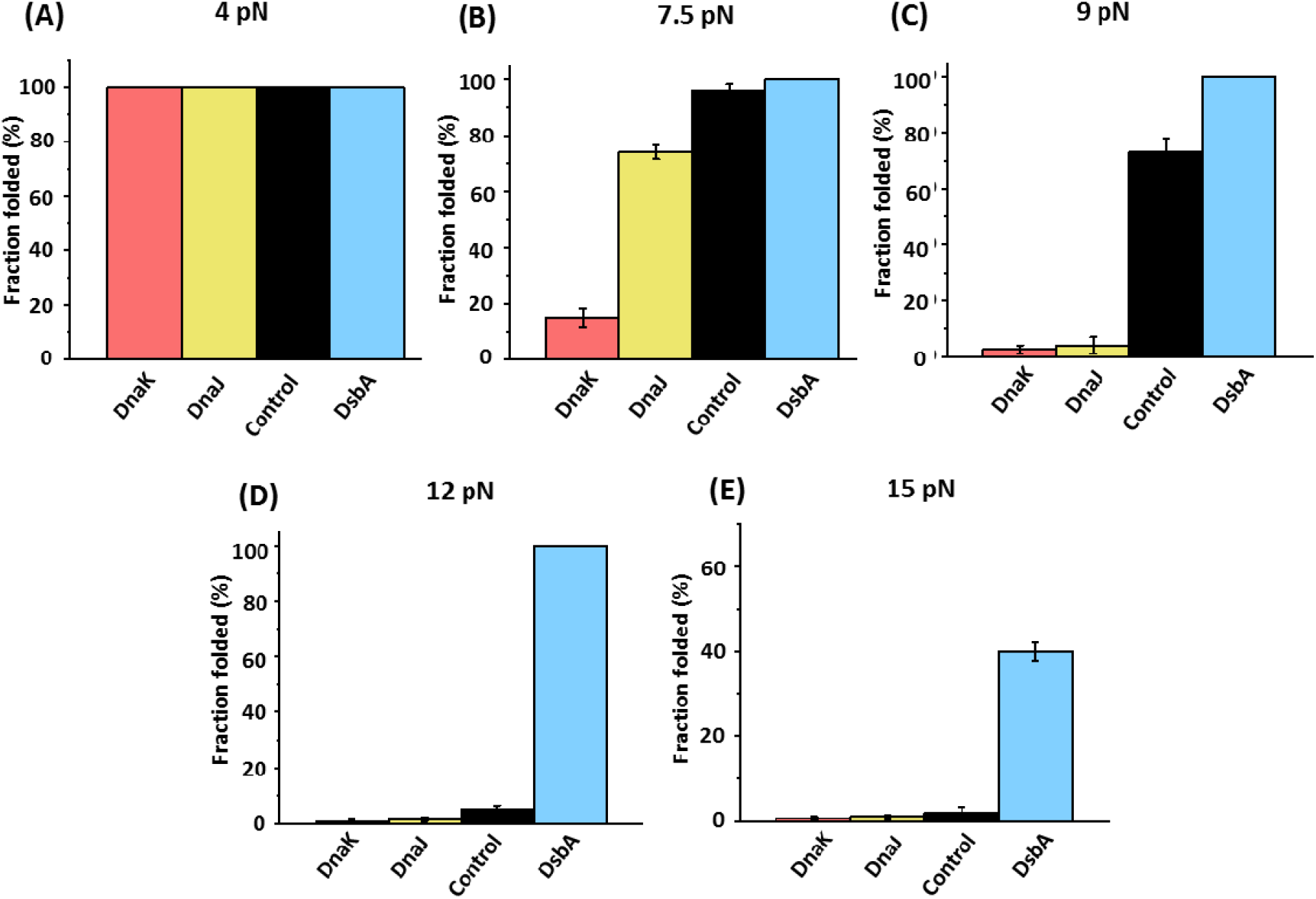
Fraction folded of talin with different chaperones at varying forces: (A) At 4 pN force, the fraction force of talin remains 100%, irrespective to chaperone presence, however, (B) and (C) the chaperones effect become more prominent at intermediate forces of 7.5 and 9 pN, where the folded fraction decreases significantly with two unfoldases. For example, at 9 pN, the fraction folded with DnaK and DnaJ are 2.6±1.5 % and 4±3.1 %, respectively; while remains 100% with DsbA due to its foldase activity, however, in control (in the absence of any chaperones), the fraction percentage decreases to only 73±5.12. (D) Since DsbA acts as a strong foldase under force, it significantly increases the talin stability by shifting the half-point force, and thus, the domain still could remain folded at 15 pN (40±2.22 %) in the presence of DsbA, while loses its ability either in control or with unfoldases.

**Supplementary Figure 13:**
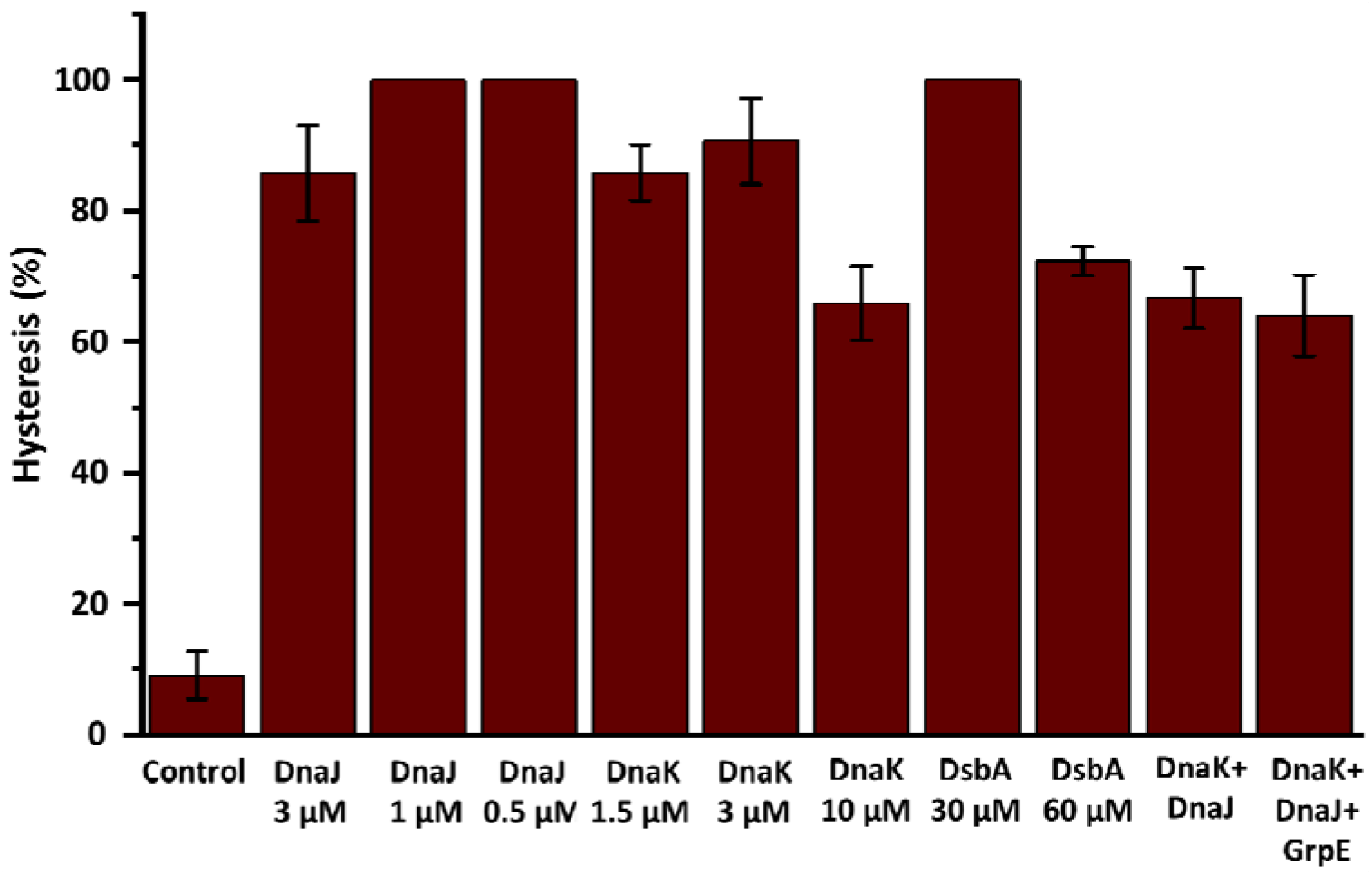
Hysteresis percentages in force-ramp experiments with different chaperones: We observed that hysteresis is substantially increased in the presence of chaperones. In the absence of chaperones or in control, the hysteresis is observed in ∼9% events, while it increases to ∼86% and ∼91% upon the addition of DnaJ and DnaK, respectively. This hysteresis could occur due to altered binding dynamics in molecular interaction where smaller hysteresis could denote rapid interaction kinetics and the larger hysteresis mean slow interaction kinetics. The mechanical unfolding of protein molecules during force-ramp experiment, is performed under non-equilibrium conditions where the complex tertiary contacts of the protein breaks and requires slower time-scale to equilibrate. This time-scale for equilibration might become further slower during the chaperone interaction, possibly resulting in the more pronounced hysteric nature in the force extension curve. Here hysteresis does not significantly affect the determination of unfolding and refolding steps as they steps are much clear and easy to detect We studied more than 50 events in each case to check the hysteresis.

### Force calibration method

In magnetic tweezers, the applied force can be empirically calibrated by several ways, depending on the construct: either by measuring the lateral fluctuation for large construct (∼10 µm) such as DNA or by relating protein unfolding extension to polymer elasticity models for smaller construct (∼50 nm) (Mora et al., Chem. Soc. Rev., 2020). Since we are using small globular protein with step size with 6-15 nm, we sought to follow the calibration method for the protein construct. Thus, we have measured the unfolding extensions against the force in PBS buffer at 25°C and fitted with the freely jointed chain (FJC) model of polymer elasticity (Fig. A) with the *Lc* value of 16.6± 0.3 nm and *Lk* of 1.1±0.2 nm. These values are in well agreement with the previous publications (Popa et al., *J. Am. Chem. Soc*., 2016). To reconcile the accuracy of the force calibration, we fitted the force at varying magnet position by exponential magnet law (Fig. B) derived by Popa et al (Popa et al., *J. Am. Chem. Soc*., 2016).

We checked the sensitivity of force calibration by imposing the deviation values of protein L polymer such as, 0.3 nm for Lc and 0.2 for Lk and observed that the effect of these deviations is within the 95% confidence level. Due to the size heterogeneity of paramagnetic beads and difference in the tether point at the bottom pole of the beads, the calibrated force has ≤10% uncertainty (Chen et al., *J. Am. Chem. Soc*., 2015; Popa et al., *J. Am. Chem. Soc*., 2016. However, the M270 paramagnetic beads have lower coefficient of variation of ∼2% than other beads of the provider, which could also ensure the negligible variation among beads. During the experiments, it is important to position the magnets by adjusting the voice coil in reproducible manner for the force application.

### Testing the accuracy of force calibration

We corroborated our force calibration method by analyzing the folding probability data of protein L molecule and then compared with the folding probability of protein L, reported by other authors who study protein nanomechanics by magnetic tweezers (Valle-Orero et al., *Angew Chem Int Ed Engl*. 2017; Valle-Orero et al., *J Phys Chem Lett*., 2017; Haldar et al., *Nat Commun*. 2017). The folding probability of protein L is highly force-dependent and we have monitored it within 4-11 pN force range. We observed that half-point force (defined as a force where FP=0.5) of protein L is ∼8 pN force and it changes significantly with 1 pN force deviation. For example, at 7 pN, the folding probability can increase by 56% (FP_7 pN_= 0.78), while decreases by 72% at 9 pN (FP_9 pN_=0.14). Thus, slight deviation in force calibration certainly affects the folding probability of protein under force, by changing the applied force on the protein tether at a particular magnet position. Notably, in our case, we found protein L exhibit half-point force at ∼8 pN force, which strongly coincides with that proposed by other groups and all the FP values are in well-agreement with their values (Fig. C). Therefore, this force calibration method has a precise detection at sub-piconewton range, allowing us to observe the force-dependent folding dynamics, which strongly claims its fiduciary force measurement by our force calibration method. Lastly, the force calibration has also been confirmed by observing the conventional B-S overstretching transition of 565 bp dsDNA at ∼66 pN, which is within the range of well-defined force standard of DNA B-S transition at 65 pN (Smith et al., *Science*, 1996).

**Supplementary Figure 14:**
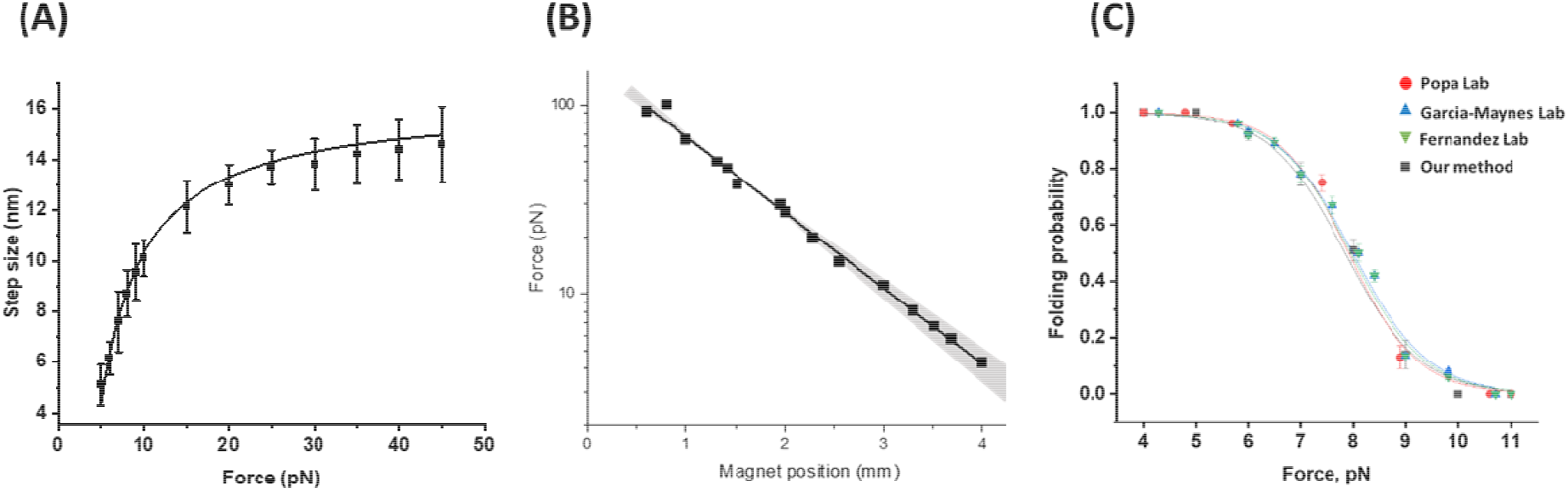
Magnet law: We calibrated the applied force using the exponential magnet law, proposed by Popa et al., JACS 2016 and demonstrates its accuracy with our instrumental set-up. **(A)** Protein L has been used as a protein force calibration and its step size or extension (∼6-15 nm) are plotted as a function of force and fitted to the freely jointed chain (FJC) model of polymer elasticity. **(B)** Accuracy of the magnet law has been reconciled by plotting the applied force against the magnet position with shaded region at 95% confidence, which is also confirmed by conventional B-S overstretching transition of 565 bp of dsDNA at 66±1.2 pN. **(C)** Furthermore, the magnet law accuracy has been tested by analyzing the folding probability of protein L with the folding probability of protein L, studied by other groups and it has been observed to overlap closely with their data.

## Notes

### Competing Interest Statement

The authors have declared no competing interest.

### Summary of Updates

Recent single-molecule studies have recognized talin as a mechanosensitive hub in focal adhesion, where its function is strongly regulated by mechanical force. For instance, at low force (below 5 pN), folded talin binds RIAM for integrin activation; whereas at high force (above 5 pN), it unfolds to activate vinculin binding for focal adhesion stabilization. Being a cytoplasmic protein, talin might interact with several cytosolic chaperones: however, the role of chaperones in talin mechanics is unknown. To address this question, we investigated the force response of a mechanically stable talin domain with a set of well-known holdase (DnaJ, DnaK, Hsp70, and Hsp40) and foldase (DnaKJE, DsbA) chaperones, using single-molecule magnetic tweezers. Our findings demonstrate that chaperone could affect adhesion proteins stability by changing their folding mechanics; while holdase chaperones reduce their unfolding force to ~6 pN, foldase chaperones shift it up to ~15 pN. Since talin is mechanically synced within 2 pN force ranges, these changes are significant in cellular condition. Furthermore, we determined the fundamental mechanism of this altered mechanical stability, where chaperones directly reshape their energy landscape: unfoldase chaperone (DnaK) decreases the unfolding barrier height from 26.8 to 21.7 kBT, while foldase chaperone (DsbA) increases it to 33.5 kBT. We reconciled our observations with eukaryotic Hsp70 and Hsp40 chaperones and observed their similar function of decreasing the talin unfolding barrier to 23.1 kBT. The quantitative mapping of this chaperone-induced talin folding landscape directly illustrates that chaperones perturb the adhesion protein stability under physiological force, thereby influencing their force-dependent interactions and adhesion dynamics.

